# Matrix stiffening promotes perinuclear clustering of mitochondria

**DOI:** 10.1101/2023.06.29.547150

**Authors:** Piyush Daga, Basil Thurakkal, Simran Rawal, Tamal Das

## Abstract

Mechanical cues from the tissue microenvironment, such as the stiffness of the extracellular matrix, modulate cellular forms and functions. As numerous studies have shown, this modulation depends on the stiffness-dependent remodeling of cytoskeletal elements. In contrast, very little is known about how the intracellular organelles such as mitochondria respond to matrix stiffness and whether their form, function, and localization change accordingly. Here, we performed an extensive quantitative characterization of mitochondrial morphology, subcellular localization, dynamics and membrane tension on soft and stiff matrices. This characterization revealed that while matrix stiffness affected all these aspects, matrix stiffening most distinctively led to an increased perinuclear clustering of mitochondria. Subsequently, we could identify the matrix stiffness-sensitive perinuclear localization of filamin as the key factor dictating this perinuclear clustering. Photo-conversion labeling and fluorescent recovery after photobleaching experiments revealed that perinuclear and peripheral mitochondrial populations differed in their motility on the soft matrix but surprisingly they did not show any difference on the stiff matrix. Finally, perinuclear mitochondrial clustering appeared to be crucial for priming human mesenchymal stem cells towards osteogenesis on the stiff matrix. Taken together, we elucidate a dependence of mitochondrial localization on matrix stiffness, which possibly enables a cell to adapt to its microenvironment.

## INTRODUCTION

The cells in our body constantly change their internal organization and adapt to various signals emanating from the extracellular environment. Mechanical cues from the extracellular matrix (ECM) elicit an intracellular tensional response that ensures tissue homeostasis. This response manifests itself in terms of cytoskeletal remodeling, altering actomyosin contractility, driving nuclear reprogramming (1), and metabolic adaptations (2, 3). Disruptions in these mechanotransduction pathways (4) have implications in cancer progression, aging, wound healing, morphogenesis, stem cell differentiation, fibrosis, and cardiovascular diseases (5-8). Till now, in the context of cellular response to mechanical cues, researchers have primarily focused on the changes in the cytoskeletal elements, including those in actin filaments, microtubules, and intermediate filaments. The cytoskeletal network, being the primary load bearing element, acts as a sink for the extracellular forces encountered by tissues. However, the participation of various organelles in cellular mechanoresponse remains mostly unknown. How these forces are eventually dissipated to organelles is not well understood. Considering that the intracellular environment is a complex web of proteins and structures where the cytoskeleton interacts with almost every organelle, cytoskeleton-generated and transmitted forces should distort organelles leading to changes in their structure and function.

Relevantly, rearrangement of intracellular structures is an energy intensive process, and the majority of the energy demand in cells is met by mitochondria which are the “powerhouses’’ of cells. Over the years, the idea of mitochondria as simply the “powerhouses’’ has expanded, and it is now clear that mitochondria act as signaling hubs, buffer calcium, aid in apoptosis and are sites of reactive oxygen species (ROS) signaling and of metabolite generation (9-11).

Moreover, our understanding of mitochondrial form has evolved dramatically over the last few decades. With improved imaging tools, it is now appreciated that mitochondria are not isolated or static. Rather they are highly dynamic and exhibit many morphologies ranging from long interconnected tubules to fragmented and swollen blobs (12). Mitochondrial dynamics, form, function, and localization are not only interlinked but are also governed by the cellular architecture (13, 14). In fact, changes in mitochondrial form or localization during biochemical stimulations, inflammations, cell division or in highly active cells like neurons, muscles and secretory cells are well characterized (15-19). In contrast, the variability in form and localization of mitochondria in response to mechanical cues, especially from the stiffness of the extracellular matrix remain poorly understood.

There are three main reasons for which it becomes imperative to study the crosstalk between matrix stiffening and mitochondrial form and function. First, there are emerging discoveries highlighting the mechanosensitivity of mitochondria to mechanical forces (20-22). In the tissue microenvironment, matrix stiffness critically influences these forces. Second, both mitochondrial dysregulation and ECM stiffening are hallmarks of cancer and aging (8, 23, 24). Finally, there exists a reciprocal relationship between cell mechanics and metabolism (3). Consequently, there are studies that have investigated the effect of matrix stiffness. For example, a recent study involving untransformed cells showed an increase in fragmented mitochondria on stiff matrix (25). On the contrary, in a study involving transformed cells, a similar phenotype was seen on soft matrix (26). Although the mitochondrial form is contradictory, both studies highlight the effect of ECM stiffness on redox homeostasis and oxidant stress responses. However, a complete picture of mitochondrial morphology, subcellular localization, and dynamics upon changes in matrix stiffness remains elusive. Here, we discover matrix stiffness as an important micro-environmental factor that distinctly decides the subcellular localization of mitochondria, in addition to its effect on morphology, motility and membrane tension.

## RESULTS

### Matrix stiffness alters morphology and subcellular localization of mitochondria

To test the effect of matrix stiffness on mitochondrial morphology and localization, we first cultured MCF-7 cells for 24 hours on collagen I-coated polyacrylamide hydrogels of elastic modulus 4 and 90 kPa, mimicking soft and stiff tissue conditions, respectively (5). We then incubated these cells with a mitochondrial potential-dependent dye Mitotracker Green, and imaged mitochondria using live confocal microscopy and super-resolution microscopy. We segmented 3D confocal images of mitochondria and quantified sphericity, which is a morphology descriptor as well as a few parameters related to mitochondrial networks, including branches per mitochondria, total branch length per mitochondria, branch junctions per mitochondria, and branch endpoints per mitochondria. On soft ECM, the number of branches, branch endpoints, and branch junctions per mitochondria were higher indicating highly networked mitochondrial structures as compared to distinct mitochondria on stiff ECM (Fig. 1a, Supp Fig. 1a-b). On stiff ECM, mitochondrial sphericity was higher indicating highly fragmented and globular mitochondria as compared to the elongated mitochondria on soft ECM (Figs. 1a-b, Supp Fig. 1a). Super-resolution images also showed highly networked mitochondria on soft ECM and toroidal mitochondria on stiff ECM (Fig. 1c). To rule out any artefact caused by live imaging and Mitotracker Green, we fixed MCF-7 cells cultured on soft and stiff ECM, stained them for the alpha subunit of mitochondrial membrane ATP Synthase or Complex V, and imaged mitochondria using confocal microscopy. Again, mitochondria on soft ECM appeared elongated and highly networked whereas mitochondria on stiff ECM appeared fragmented and toroid-shaped (Fig. 1d, Supp Fig. 1c). These results showed that imaging conditions and specific stains did not affect mitochondrial morphology and network properties on soft and stiff ECM. Along with the stiffness-dependent changes in the mitochondrial morphology, we also observed distinct mitochondrial localization patterns. Mitochondria on soft ECM were homogeneously distributed across the cell area, whereas mitochondria on stiff ECM were predominantly clustered near the nucleus (the perinuclear space) (Figs. 1a, d). Since cells are reported to spread more on stiff ECM (5), we decided to check whether mitochondrial localization was dependent on cell spreading. To this end, we plotted the distribution of cytoplasm and mitochondria (localization distribution function) as a function of distance from the nucleus and further quantified the percent of mitochondria localized in a defined perinuclear radius (5 µm). The distribution of the cytoplasm was similar on either stiffness condition, but the mitochondrial distribution showed a distinct peak in the perinuclear zone on stiff ECM, representing the perinuclear clustering of mitochondria (Fig. 1e), confirming that mitochondrial localization was not dependent on cell spreading. Taken together, these results showed that matrix stiffening creates distinct mitochondrial populations - homogeneously distributed, elongated, and networked tubules on soft ECM and fragmented perinuclear clusters on stiff ECM. However, the effect of matrix stiffness on mitochondrial localization appeared more distinct than on mitochondrial morphology (Fig. 1g). On stiff ECM, mitochondria appeared to be distinctly clustered around the nucleus, and a major region of the cytoplasm lacked mitochondria (Fig. 1e). Hence, subsequently we tested whether stiffness-sensitive mitochondrial localization depended on cell-ECM adhesions (Figs. 1f-g), probed the motility of perinuclear versus peripheral mitochondrial populations (Fig. 2), asked what caused the differential localization (Fig. 3), measured the mitochondrial membrane tension (Fig. 4), and finally, elucidated the physiological relevance of perinuclear clustering of mitochondria on stiff ECM to stiffness-mediated change in the stem-cell fate (Fig. 5).

**Figure 1:**
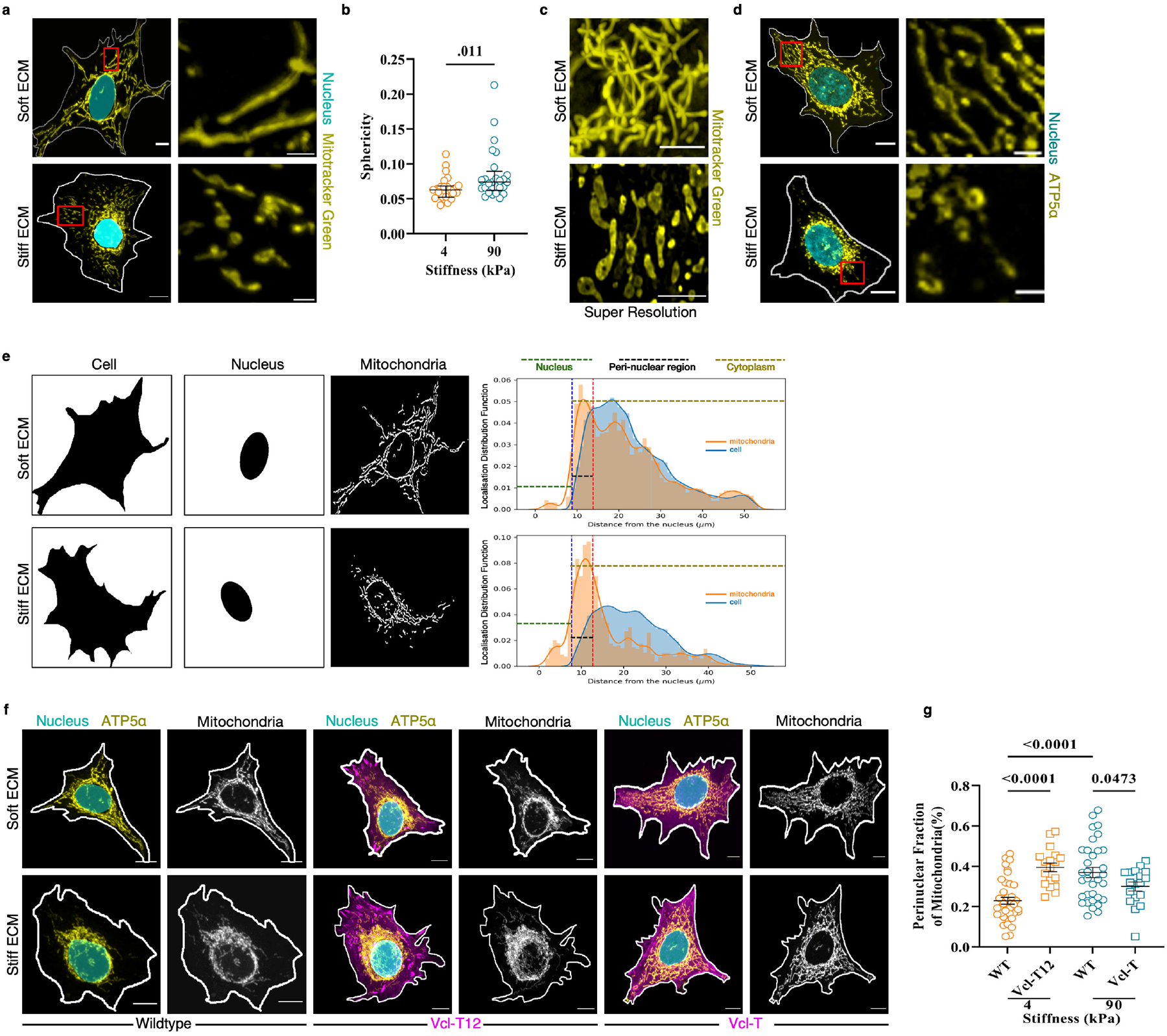
Matrix stiffness alters morphology and cellular localization of mitochondria. **a**.3D stacked confocal images of MCF-7 cells cultured on soft (top panel) and stiff (bottom panel) ECM showing mitochondria stained with Mitotracker Green. From left to right: Merged images showing nuclei in cyan and mitochondria in yellow followed by a magnified view of mitochondria in the red box (scale bar: 2 μm). **a**. Scatter dot plot showing weighted sphericity of mitochondria per cell cultured on soft (4 kPa) and stiff (90 kPa) ECM. n=24 FOVs (4 kPa), 25 FOVs (90 kPa). Each FOV (field of view) contains 10-15 cells. **b**. Representative super-resolution image of MCF-7 cells cultured on soft (top panel) and stiff (bottom panel) ECM showing mitochondria (yellow) stained with Mitotracker Green. **c**. Immunofluorescence confocal images of MCF-7 cells cultured on soft (top panel) and stiff (bottom panel) ECM showing nuclei stained with DAPI and mitochondria stained with ATP5α. From left to right: Merged images showing nuclei in cyan and mitochondria in yellow followed by a magnified view of mitochondria in the red box (scale bar: 2 μm). **d**. Representative binarized images of cell body, nucleus and mitochondria on soft ECM (top panel) and stiff ECM (bottom panel). From left to right: Binarized images showing area of a single cell (in black) followed by nuclear area (in black) and mitochondria positive pixels (in white), graphs showing distribution of cytoplasm (blue) and mitochondria (orange) plotted as a function of distance from the nucleus (localization distribution function). The nuclear, perinuclear and cytoplasmic zones are indicated on the graphs. **e**. 3D stacked confocal images of MCF-7 cells cultured on soft (top panel) and stiff (bottom panel) ECM under untransfected (wildtype) and transfected conditions showing mitochondria stained with ATP5α. From left to right: Merged image of MCF-7 wildtype cells showing nuclei in cyan and mitochondria in yellow followed by mitochondria in grayscale highlighting differences in localization, merged image of MCF-7 cells transfected with Vcl-T12 (magenta) showing nuclei in cyan and mitochondria in yellow followed by mitochondria in grayscale showing perinuclear clustering (also shown in Supp Fig. 2b), merged image of MCF-7 cells transfected with Vcl-T (magenta) showing nuclei in cyan and mitochondria in yellow followed by mitochondria in grayscale showing homogenous distribution (also shown in Supp Fig. 2c). **f**. Scatter dot plot showing fraction of mitochondria present in the perinuclear zone in the wild-type (WT) and Vcl-T12 transfected cells cultured on soft ECM (4 kPa) and wild-type (WT) and Vcl-T transfected cells cultured on stiff ECM (90 kPa). n= 38 (WT, 4 kPa), 18 (Vcl-T12, 4 kPa), 39 (WT, 90 kPa), 18 (Vcl-T, 90 kPa) cells. Statistical significance was calculated using two-tailed Mann-Whitney test (**b**) or Brown-Forsythe Anova test with Welch’s correction (**g**). Data are given as median and interquartile range (**b**) or mean ± s.e.m (**g**) taken over two independent replicates. P-values are shown in the graphs. White lines denote the cell boundary in (**a**,**d**,**f**) and cyan ovals depict cell nuclei in (**a**,**e**), both drawn manually using DIC images (for wild-type cells) or GFP fluorescence (for transfected cells) that were acquired simultaneously with fluorescence images. Scale bar: 10 μm. Soft ECM: 4 kPa; Stiff ECM: 90 kPa

### Stiffness-sensitive mitochondrial localization, but not morphology, depends on integrin signaling

MCF-7 cells extended protrusions on soft and stiff ECM indicating mechanical engagement on either stiffness condition. Furthermore, these cells showed increased spreading and distinct actin stress fibers on stiff ECM (Supp Fig 2a), indicating integrin-based mechano-sensing of ECM stiffness. Hence, to further understand the effect of ECM stiffness on the subcellular localization of mitochondria, we targeted integrin-mediated mechano-signaling, which connects the cytoskeleton to ECM forming a mechanical continuity (4). Vinculin is an integral part of integrin signaling, serving as a molecular link between talin and the actin cytoskeleton (27). Vinculin binds to talin via its head domain and to filamentous-actin (F-actin) through its tail domain. However, vinculin exists in an auto-inhibited state where intramolecular interactions between head and tail domains mask ligand binding sites. Under mechanical tension, vinculin gets activated leading to conformational changes that dissociate the head-tail interaction and expose the binding sites (28). We asked whether mitochondrial morphology and localization might be sensitive to Vinculin-mediated integrin signaling. To this end, we used two vinculin plasmids: a constitutively active form of Vinculin labeled with green fluorescent protein, where the head-tail interaction was abolished (called Vcl-T12) and Vinculin tail domain labeled with green fluorescent protein, which only binds to actin but not talin (called Vcl-T) (28). Vcl-T binds to the barbed ends of F-actin and hence prevents actin polymerization, thereby having a dominant negative effect on the interaction between endogenous vinculin and F-actin. (29, 30). We first enhanced integrin mechanosignaling by transient expression of Vcl-T12 in MCF-7 cells cultured on soft ECM. Cells positive for Vcl-T12 showed distinct actin filaments in contrast to diffused actin structures seen in wildtype cells on soft ECM (Supp Fig. 2a). Conversely, we abrogated integrin signaling by transient expression of Vcl-T in MCF-7 cells cultured on stiff ECM. Cells positive for Vcl-T had diffused actin structures in contrast to distinct stress fibers seen in wildtype cells on stiff ECM (Supp Fig. 2a). Mitochondrial localization appeared very sensitive to mutant Vinculin-changes. On soft ECM, cells positive for Vcl-T12 showed increased perinuclear clustering of mitochondria in contrast to the homogeneously distributed mitochondria seen in the wild-type cells (Figs. 1f-g, Supp Fig. 2b). On stiff ECM, cells positive for Vcl-T showed homogenous distribution of mitochondria in contrast to the perinuclear clustering seen in the wild-type cells (Figs. 1f-g, Supp Fig. 2c). However, Vinculin mutants did not have any significant effect on mitochondrial morphology or network properties on either ECM stiffness (Supp Fig. 1d). Taken together, these results clearly showed that mitochondrial positioning within a cell critically depends on integrin-mediated sensing of ECM stiffness.

### Matrix stiffness generates distinctly dynamic mitochondrial populations

We subsequently characterized the dynamical properties of mitochondria on soft and stiff ECM and asked whether there could be any localization-specific differences among mitochondria on soft and stiff ECM. To this end, we stably expressed MitoDendra2, a photo-switchable fluorescent protein construct targeted to the mitochondrial matrix (31), in MCF-7 cells. We cultured these cells on soft and stiff ECM. We then imaged mitochondrial dynamics by 2D time lapse confocal microscopy and used an automated toolkit for mitochondrial segmentation and tracking, namely Mitometer (32), to analyze motility features such as the distance, speed, displacement, and velocity. This analysis showed that all of the motility features of mitochondria were higher on stiff ECM than on soft ECM (Supp Fig. 3a), implying mitochondria in general are more mobile on stiff ECM than on soft ECM. Next, considering the effect of ECM stiffness on mitochondrial localization, we asked whether this difference arose due to differences in the motility of specific mitochondrial subpopulations or it might be general and location-independent. To test these alternative possibilities, we photoconverted a population of MitoDendra2 expressing mitochondria at two locations each: one in the perinuclear (PN) region and one away from the nucleus, in the peripheral (PL) region. We then monitored the diffusion of the photo-converted molecules using time lapse imaging techniques. Analysis of photo-converted mitochondria showed that on soft ECM, the velocity, displacement, distance, and speed of PL mitochondria were higher than PN mitochondria, while on stiff ECM the same parameters were comparable between PN and PL mitochondria. (Figs. 2a-b, Supp Fig. 3b, Supp Videos 1-2). In short, it was surprising that soft ECM created two mitochondrial populations that were different in their motilities.

**Figure 2:**
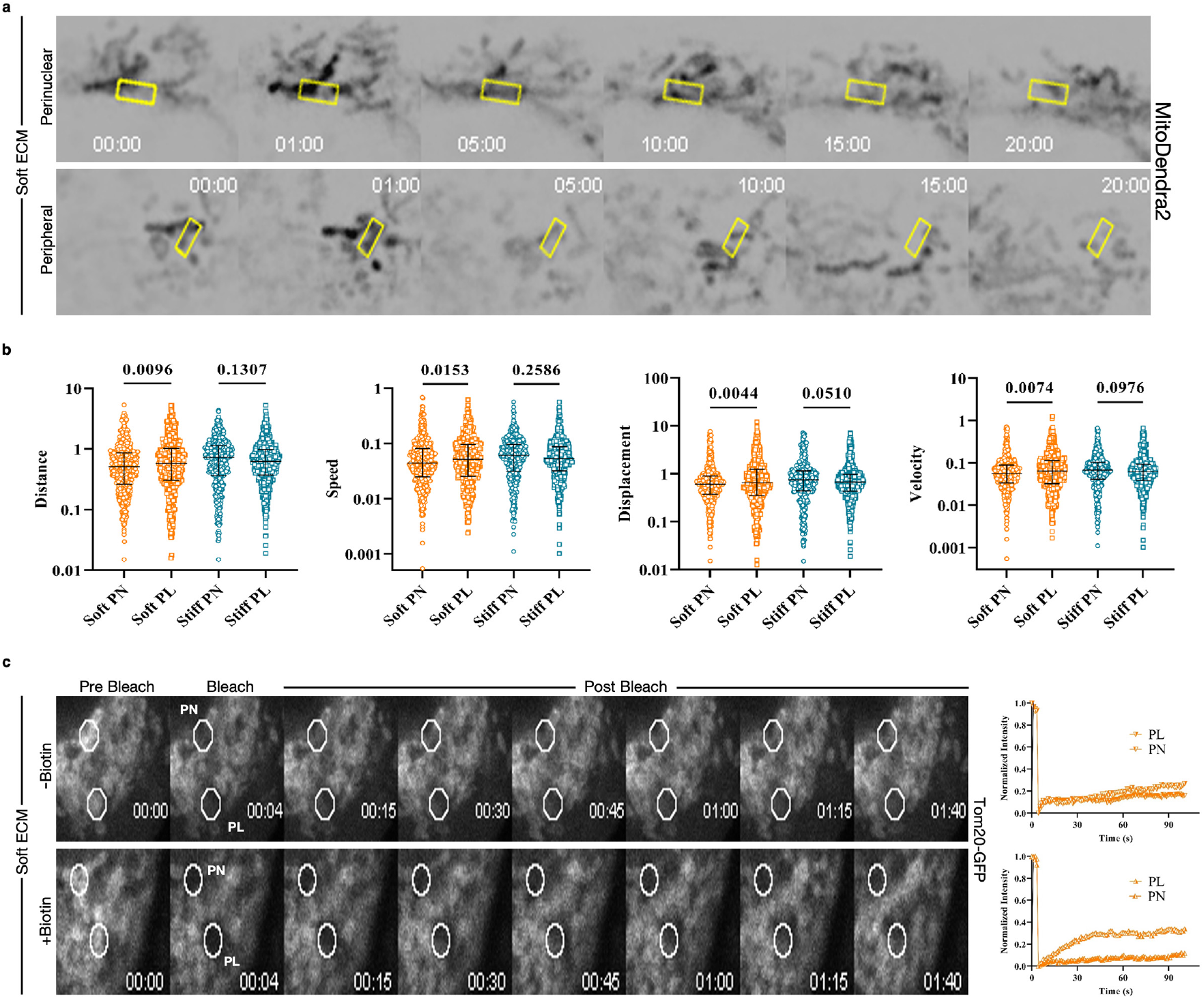
Matrix stiffness generates distinctly dynamic mitochondrial populations. **a**. Montage showing mitochondrial motility at perinuclear (PN) and peripheral (PL) regions of MitoDendra2 expressing MCF-7 cells cultured on soft ECM. Yellow boxes represent ROIs where Dendra2 expressing mitochondria were photoconverted (t=00:00) and subsequently the diffusion of the photoconverted molecules was tracked for 20 mins (also see Supp Video 1). **b**. Scatter dot plots showing motility metrics: distance (μm), speed (μm s^-1^), displacement (μm) and velocity (μm s^-1^) of photo converted mitochondria in PN and PL regions of MitoDendra2 MCF-7 cells cultured on soft and stiff ECM. n= 546 (soft PN), 594 (soft PL), 469 (stiff PN), 675 (Stiff PL) distinct mitochondrial tracks. Statistical significance was calculated using Kruskal-Wallis nonparametric test with uncorrected Dunn’s test. Data are given as median and interquartile range taken over two independent replicates. P-values are shown in the graphs. The Y-axes of plots showing distance, speed, displacement, and velocity are on the “log 10” scale. **c**. Montage showing fluorescence recovery after photobleaching in perinuclear (PN) and peripheral (PL) regions of MCF-7 cells transfected with mCh-KIF5C and Tom20-GFP and cultured on soft ECM without Biotin (top panel) and with Biotin (bottom panel). White circles represent photo-bleached regions. From left to right: GFP Fluorescence in PN and PL ROIs (white circles) captured at pre-bleach time point (t=00:00) followed by bleaching (t=00:04) and subsequent recovery tracked for 1 min 40 s, FRAP curves showing fluorescence recovery of Tom20-GFP at the indicated ROIs (also see Supp Video 3-4). ROI: region of interest. Soft ECM: 4 kPa; Stiff ECM: 90 kPa

To further test whether the mitochondrial localization was indeed influencing the mitochondrial motility, we used the reversible association with motor proteins (RAMP) system to manipulate mitochondrial positioning within the cell (33). In particular, we used the retrograde motor protein Kif5C1 fused to streptavidin and mCherry (mCh-KIF5C) and mitochondria outer membrane protein TOM20 fused to streptavidin binding protein (SBP) and GFP (Tom20-GFP). The SBP-Streptavidin interaction drives all the mitochondria towards the minus end of microtubules, near the nucleus. Importantly, this interaction is reversible as addition of biotin outcompetes the SBP-Streptavidin interaction and allows restoration of normal mitochondrial distribution (33). Therefore, we used this system to localize a major population of mitochondria near the nucleus on soft ECM and studied whether this imposed perinuclear localization would perturb the motility. To this end, we transiently co-expressed mCh-KIF5C and Tom20-GFP to promote perinuclear mitochondrial localization on soft ECM and then studied mitochondrial motility by measuring the fluorescent recovery of the photobleached GFP. We selected two sets of locations for photobleaching : one in the perinuclear (PN) region and one away from the nucleus (peripheral or PL). Cells positive for both plasmid constructs indeed had increased perinuclear mitochondria (Supp Fig. 3d). Subsequently, fluorescent recovery after photobleaching (FRAP) analysis showed that the mobile fraction in PN and PL mitochondria was comparable (Fig. 2c, Supp Fig. 3e, Supp Video 3). However, upon addition of biotin which reverted mitochondria to its original homogenous distribution, the mobile fraction of PL mitochondria was greater than PN mitochondria (Fig. 2c, Supp Fig. 3e, Supp Video 4), reiterating our previous observation that on soft ECM, PL mitochondria are more mobile (Fig. 2b). As expected, on stiff ECM, where mitochondria are anyways predominantly localized in the perinuclear region, the mobile fractions of PL and PN mitochondria were comparable before and after biotin treatment (Supp Fig. 3e).

These results show that specific perinuclear clustering of mitochondria on stiff ECM abolishes differences in motility that were seen in homogeneously distributed mitochondria on soft ECM. In fact, both correlation experiments (with MitoDendra2) and causation experiments (with RAMP system) converged to the same conclusion that soft ECM creates two mitochondrial populations that differ in their motilities. Upon perinuclear clustering, this difference gets abrogated. It is possible that the motility difference in PN and PL mitochondrial populations on soft ECM arises due to the difference in the intracellular volume constraint. The perinuclear region being relatively more organelle-filled than the peripheral region, should impose higher restrictions to mitochondrial motility. But importantly the fact that this difference gets abrogated on stiff ECM indicates that there must be some intracellular elements that actively interact with mitochondria in the perinuclear space on stiff ECM.

### Perinuclear accumulation of filamin alters mitochondrial localization

To seek out these elements, we next asked what factors determined the differential localization of mitochondria on soft and stiff ECM. It is known that ECM stiffness alters the cellular localization of several force-sensitive cytoskeleton-related proteins (34). Recently, we have shown that ECM stiffness sensitive differential localization of an actin crosslinker protein, filamin (filamin A or FLNA), plays a critical role in epithelial defense against cancer (35), where perinuclear filamin molecules co-localize with perinuclear actin cytoskeleton on stiff ECM. In this current study, we also found that while total filamin levels on soft and stiff ECM were comparable as quantified by immunostaining and microscopy, a significant fraction of filamin was localized in the perinuclear region on stiff ECM (Figs. 3a-b, Supp Fig. 4a) corroborating our previous results (35). Here we speculated that filamin could be an interesting candidate with respect to stiffness-sensitive mitochondrial localization, considering the intricate association of actin structures with mitochondrial dynamics (36-41) and the reported interaction between filamin and mitochondrial dynamics related proteins (42). We, therefore, hypothesized that stiffness-sensitive differential localization of filamin might be responsible for the stiffness-sensitive differential localization of mitochondria. To test this hypothesis, we transiently overexpressed filamin, labeled with fluorescent protein mApple (called mAppFLNA hereafter), in MCF-7 cells cultured on soft ECM. Filamin overexpression increased filamin distribution throughout the cell, especially in the perinuclear region ((35), Fig. 3c). Interestingly, cells overexpressing filamin showed increased perinuclear mitochondrial clustering even on soft ECM in contrast to the homogenous distribution seen in the wildtype cells. However, filamin overexpression did not further affect perinuclear mitochondrial clustering on stiff ECM (Figs. 3d-f, Supp Figs. 4b-c), which were already clustered around the nucleus. We next tested the effect of disruption of perinuclear filamin on mitochondrial localization. Considering that refilinB-filamin interaction is important for localizing filamin to the perinuclear cytoskeleton (35, 43), we disrupted this interaction and reduced the perinuclear filamin localization on stiff ECM by transient expression of a dominant negative filamin mutant (called dnFLNA hereafter) (Fig. 3c). dnFLNA carries only four filamin repeats and is known to have a dominant negative effect on the interaction between endogenous filamin and refilinB (43). Cells expressing dnFLNA (labeled with mApple) indeed showed homogeneous distribution of mitochondria on stiff ECM in contrast to the perinuclear clustering seen in the wildtype cells. Interestingly, dnFLNA expression did not affect mitochondrial localization on soft ECM (Figs. 3d-f, Supp Figs. 4b, d). Taken together, these results showed that ECM stiffening-induced perinuclear enrichment of filamin affected mitochondrial localization, and perinuclear filamin aggregation is critical for perinuclear clustering of mitochondria.

**Figure 3:**
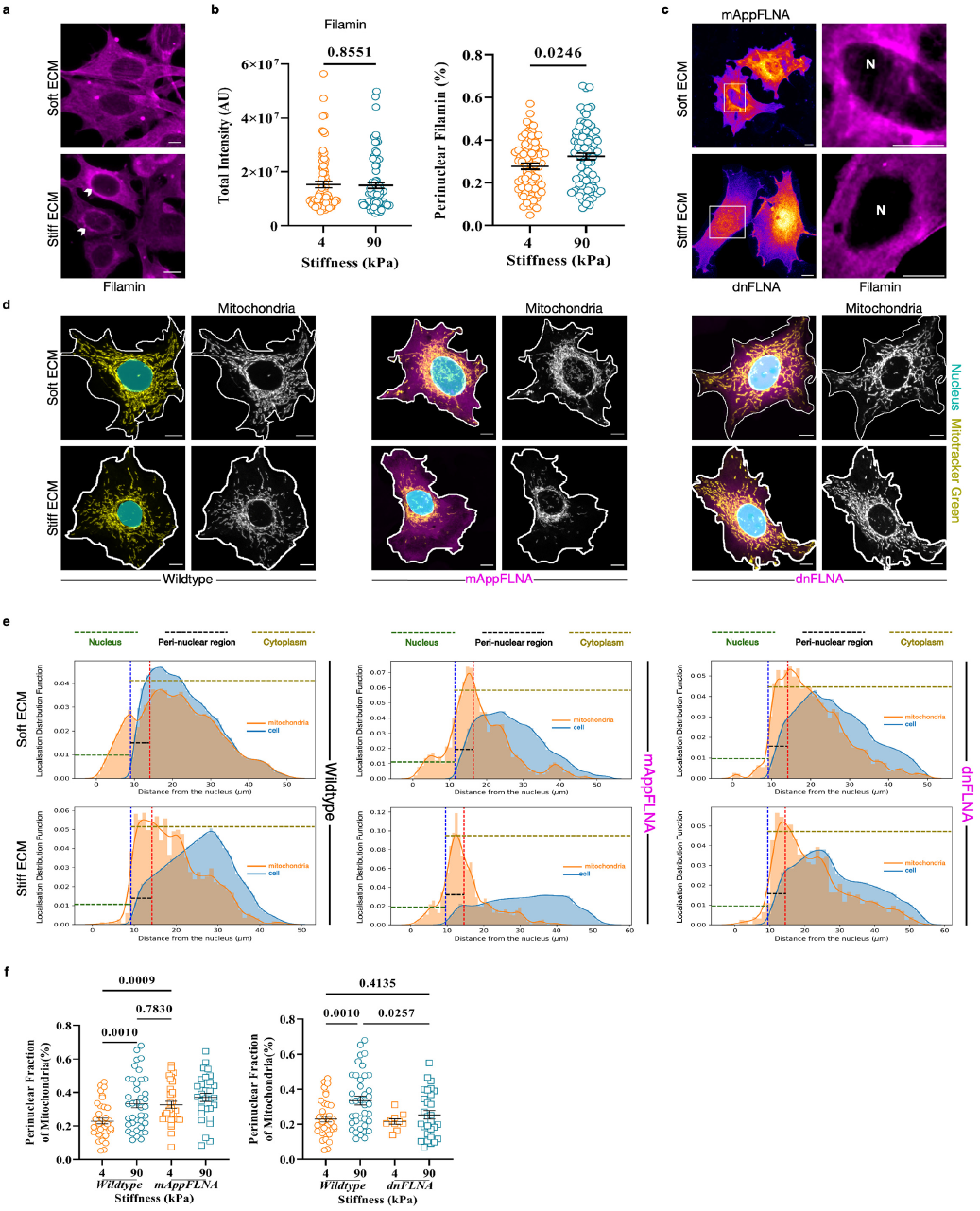
Perinuclear accumulation of filamin alters mitochondrial localization. **a**. Immunofluorescence images of MCF-7 cells stained for filamin cultured on soft (top panel) and stiff (bottom panel) ECM. Arrowheads show perinuclear enrichment of filamin on stiff ECM. **b**. Scatter dot plots depicting total filamin protein levels (total intensity) and fraction of filamin in the perinuclear region (perinuclear filamin %) on soft (4 kPa) and stiff (90 kPa) ECM. n= 78 (4 kPa), 83 (90 kPa) cells. **c**. Top panel: Confocal images of MCF-7 cells transiently expressing mAppFLNA cultured on soft ECM followed by a magnified view of the white box showing perinuclear enrichment of filamin (in magenta). Bottom panel: Confocal images of MCF-7 cells transiently expressing dnFLNA cultured on stiff ECM followed by a magnified view of the white box showing reduced perinuclear localization of endogenous filamin (in magenta). N: nucleus. **d**. 3D stacked confocal images of MCF-7 cells cultured on soft (top panel) and stiff (bottom panel) ECM under untransfected (wild-type) and transfected conditions showing mitochondria stained with Mitotracker Green. From left to right: Merged image of MCF-7 wild-type cells showing nuclei in cyan and mitochondria in yellow followed by mitochondria in grayscale highlighting differences in localization, merged image of MCF-7 cells transfected with mAppFLNA (magenta) showing nuclei in cyan and mitochondria in yellow followed by mitochondria in grayscale showing perinuclear clustering, merged image of MCF-7 cells transfected with dnFLNA (magenta) showing nuclei in cyan and mitochondria in yellow followed by mitochondria in grayscale showing homogenous distribution. White lines denote the cell boundary and cyan ovals depict cell nuclei, both drawn manually using DIC images (for wild-type cells) or mApple fluorescence (for transfected cells) that were acquired simultaneously with other fluorescence images. **e**. Graphs showing distribution of cytoplasm (in blue) and mitochondria (in orange) in cells shown in (**d**) plotted as a function of distance from the nucleus. From left to right: wild-type cells, mAppFLNA transfected cells and dnFLNA transfected cells cultured on soft ECM (top panels) and stiff ECM (bottom panels). The nuclear, perinuclear and cytoplasmic zones are indicated on the graphs. The respective binarized images are shown in Supp Figs. 4b-d. **f**. Scatter dot plots showing fraction of mitochondria present in the perinuclear zone in wild-type and mAppFLNA (left graph) or dnFLNA (right graph) transfected cells cultured on soft (4 kPa) and stiff (90 kPa) ECM. n= 38 (wild-type, 4 kPa), 39 (wild-type, 90 kPa), 32 (mAppFLNA, 4 kPa), 30 (mAppFLNA, 90 kPa), 10 (dnFLNA, 4 kPa), 29 (dnFLNA, 90 kPa) cells. Statistical significance was calculated using unpaired t-test with Welch’s correction (two-tailed) (**b**) or Brown-Forsythe Anova test with Welch’s correction (**f**). Data are given as mean ± s.e.m (**b**,**f**) taken over two independent replicates. P-values are shown in the graphs. Scale bar: 10 μm. Soft ECM: 4 kPa; Stiff ECM: 90 kPa

### Mitochondrial membrane tension is sensitive to matrix stiffness but not to mitochondrial localization

Considering that filamin is an important cytoskeleton-associated protein, and cellular organelles are known to experience physical cues through their association with the cytoskeleton (13, 14), we next asked whether the filamin-dependent mitochondrial localization would also influence physical properties of mitochondria. Relevantly, mitochondria are tubular structures and bear tension during force loading (22, 40). Since mitochondrial membrane tension plays a role in mitochondrial dynamics (20), we examined how stiffness mediated changes in mitochondrial morphology and localization would affect mitochondrial membrane tension. To measure tension, we used Mito Flipper-TR (44), a mitochondrial-targeted mechanosensitive FliptR probe that reports changes in membrane tension through its fluorescence lifetime. We confirmed the mitochondrial localization of the flipper probe by co-staining with an established mitochondrial marker (Mitotracker Red) (Supp Fig. 5a). Upon Fluorescence Lifetime Imaging Microscopy (FLIM) analysis, we observed that mitochondrial membrane tension as indicated by fluorescence lifetimes of Mito Flipper-TR probe was lowered on stiff ECM as compared to soft ECM (Figs. 4a-b). Fluorescence lifetimes in perinuclear and peripheral mitochondrial populations on either stiffness condition were comparable (Fig. 4b). To further test the effect of mitochondrial localization on mitochondrial membrane tension, we measured membrane tension in mAppFLNA expressing cells on soft ECM and in dnFLNA expressing cells on stiff ECM (Fig. 4c). Perinuclear aggregation of mitochondria by mAppFLNA did not alter fluorescence lifetimes in perinuclear or peripheral mitochondrial populations when compared to the wild-type (untransfected) cells on soft ECM. Conversely, disruption of perinuclear mitochondrial clusters by dnFLNA did not alter fluorescence lifetimes in perinuclear or peripheral mitochondrial populations when compared to the wild-type (untransfected) cells on stiff ECM (Fig. 4d-e). Taken together, these results showed that the subcellular localization of mitochondria has no effect on membrane tension. However, mitochondria adapt to a stiffer microenvironment by preferring to exist as fragments having low membrane tension.

**Figure 4:**
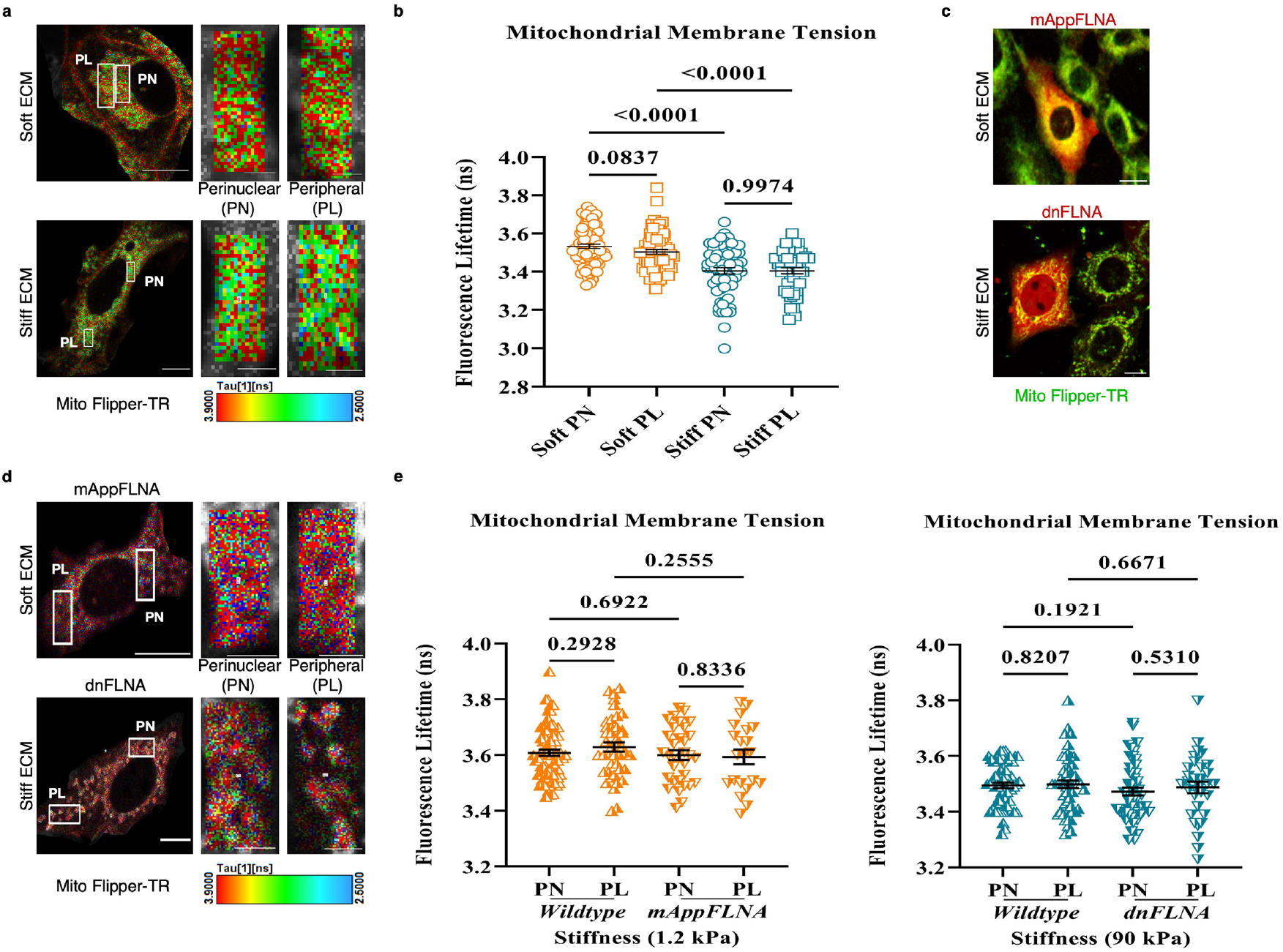
Mitochondrial membrane tension is sensitive to matrix stiffness but not to mitochondrial localization. **a**. FLIM images of MCF-7 cells cultured on soft (top panel) and stiff (bottom panel) ECM showing mitochondria stained with Mito Flipper-TR. White boxes represent ROIs drawn in the perinuclear (PN) and peripheral (PL) regions that were used to calculate fluorescence lifetimes. Magnified view of ROIs (scale bar: 2 μm) shows higher fluorescence lifetimes on soft ECM. Scale bar:10 μm. **b**. Scatter dot plot showing average fluorescence lifetimes calculated in PN and PL regions of cells cultured on soft and stiff ECM. n= 82 (soft PN), 68 (soft PL), 67 (stiff PN), 50 (stiff PL) ROIs from cells (83 cells on soft ECM, 76 cells on stiff ECM). **c**. Confocal images of MCF-7 cells transfected with mAppFLNA (in red) and cultured on soft ECM (top panel) and with dnFLNA (in red) and cultured on stiff ECM (bottom panel) showing mitochondria (in green) stained with Mito Flipper-TR. **d**. FLIM images of MCF-7 cells transfected with mAppFLNA and cultured on soft ECM (top panel); and with dnFLNA and cultured on stiff ECM (bottom panel) showing mitochondria stained with Mito Flipper-TR. White boxes represent ROIs drawn in the perinuclear (PN) and peripheral (PL) regions that were used to calculate fluorescence lifetimes. Magnified views of ROIs (scale bar: 5 μm) are shown for comparison. Scale bar:20 μm. **e**. Scatter dot plot showing average fluorescence lifetimes calculated in PN and PL regions of wild type (untransfected) and mAppFLNA transfected cells cultured on soft ECM (left graph); and wildtype (untransfected) and dnFLNA transfected cells cultured on stiff ECM (right graph). n=76 (wild type PN), 42 (wild type PL), 33 (mAppFLNA PN), 21 (mAppFLNA PL) ROIs from cells (42 wild type cells and 21 mAppFLNA transfected cells on 1.2 kPa); n=62 (wild type PN), 54 (wild type PL), 53 (dnFLNA PN), 32 (dnFLNA PL) ROIs from cells (47 wild type cells and 31 dnFLNA transfected cells on 90 kPa). Statistical significance was calculated using unpaired t-test with Welch’s correction (two-tailed) (**b**,**e**). Each data point represents the average lifetime in each ROI. Data are mean ± s.e.m (**b**,**e**) taken over two independent replicates. P-values are shown in the graphs. ROI: region of interest. Soft ECM: 1.2 kPa; Stiff ECM: 90 kPa

### Perinuclear mitochondrial clustering primes mesenchymal stem cells towards osteogenesis

Finally, we asked what might be the physiological relevance of perinuclear mitochondrial clustering on stiff ECM. It is well known that culturing human mesenchymal stem cells (hMSCs) on stiff ECM promotes osteogenesis, generally marked by increased nuclear expression of an osteogenesis-related protein RUNX2 (5). In this respect, at first, examining the mitochondrial morphology and localization in hMSCs, we obtained similar results as before in MCF-7 cells. Mitochondria in hMSCs cultured on soft ECM were highly networked, elongated and homogeneously distributed whereas mitochondria in hMSCs cultured on stiff ECM were fragmented and formed perinuclear clusters (Figs. 5a-d, Supp Figs. 6a-b). Interestingly, we also observed that transient expression of mAppFLNA in MSCs on soft ECM led to mitochondrial clustering near the nucleus, whereas perinuclear filamin disruption using dnFLNA on stiff ECM led to homogeneous distribution of mitochondria (Figs. 5c-d, Supp Fig. 6b), further extending the generality of our findings. Given these findings, we hypothesized that if mitochondrial localization on stiff ECM indeed influenced osteogenic differentiation of hMSCs, perturbing the former by expressing dnFLNA would attenuate osteogenesis on stiff ECM. To test this hypothesis, we cultured hMSCs in normal media conditions (undifferentiated state) on soft or stiff ECM for 24 hours. Upon immunostaining for RUNX2, we observed that this osteogenic marker was lowly expressed on soft ECM, whereas the expression was detectable in the cell nuclei on stiff ECM (Supp Fig. 6c). We then disrupted perinuclear mitochondrial clustering on stiff ECM via transient expression of dnFLNA and observed that the nuclear signal of osteogenic marker RUNX2 reduced significantly when compared to wildtype cells (Fig. 5e, Supp Fig. 6d). dnFLNA mediated disruption affected mitochondrial localization and by extension reduced nuclear localization of RUNX2 on stiff ECM. Taken together, these results show that perinuclear mitochondrial clustering on a stiffened matrix might be crucial for priming stem cells towards osteogenesis (Fig. 5f).

**Figure 5:**
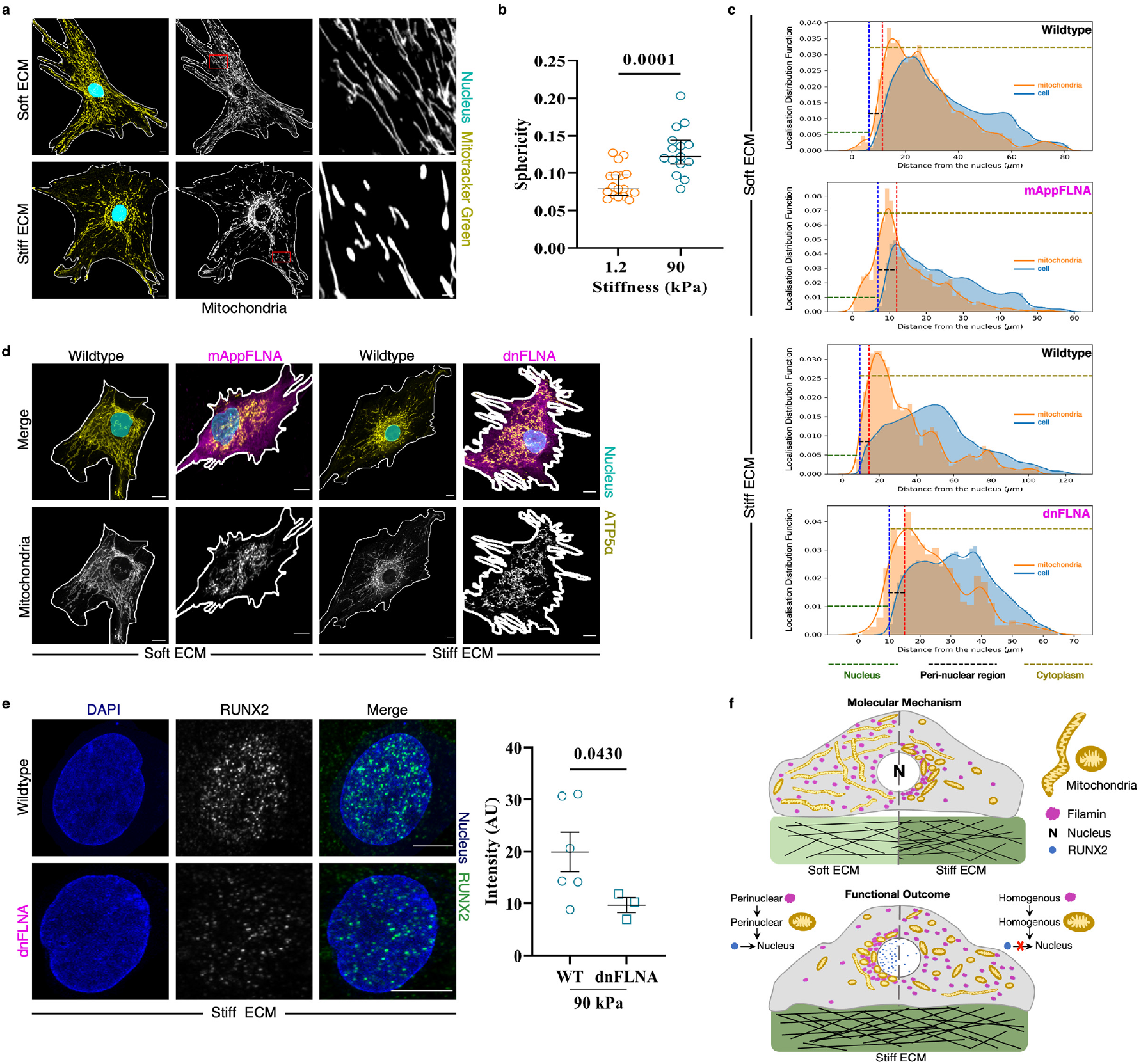
Perinuclear mitochondrial clustering prime mesenchymal stem cells towards osteogenesis. **a**. 3D stacked confocal images of hMSCs cultured on soft (top panel) and stiff (bottom panel) ECM showing mitochondria stained with Mitotracker Green. From left to right: Merged images showing nuclei in cyan and mitochondria in yellow, grayscale images of mitochondria showing differences in localization, magnified view of the red boxes (scale bar: 2 μm) showing differences in mitochondrial morphology. **b**. Scatter dot plot showing weighted sphericity of mitochondria per cell cultured on soft (1.2 kPa) and stiff (90 kPa) ECM. n=18 FOVs (1.2 kPa), 15 FOVs (90 kPa) taken over two independent replicates. Each FOV (field of view) contains 3-5 cells. **c**. Graphs showing distribution of cytoplasm (in blue) and mitochondria (in orange) in cells shown in (**d**) plotted as a function of distance from the nucleus. From top to bottom: wild-type and mAppFLNA transfected hMSCs cultured on soft ECM, wild-type and dnFLNA transfected hMSCs cultured on stiff ECM. The nuclear, perinuclear and cytoplasmic zones are indicated on the graphs. The respective binarized images are shown in Supp Fig. 6b. **d**. 3D stacked confocal images of hMSCs transfected with mAppFLNA and cultured on soft ECM (left panels); and with dnFLNA and cultured on stiff ECM (right panels) showing mitochondria immunostained with ATP5α.From top to bottom: Merged images showing nuclei in cyan and mitochondria in yellow followed by grayscale images of mitochondria showing differences in localization (also represented in (**c**)). **e**. Immunofluorescence images of hMSCs cultured on stiff ECM under untransfected (wild-type, top panel) and transfected (dnFLNA, bottom panel) conditions and stained with DAPI and RUNX2 (also see Supp Fig. 6d). From left to right: Confocal images showing nuclei stained with DAPI, grayscale images of RUNX2 expression in nuclei, merged images of nuclei in blue and RUNX2 in green, scatter dot plot showing average intensity of RUNX2 in nuclei of WT (wild-type) and dnFLNA transfected cells on 90 kPa. n= 6 (WT), 3 (dnFLNA) cells. **f**. Schematic summarizing the key findings: Matrix stiffening affects mitochondrial localization. Mitochondria on soft ECM are homogeneously distributed across the cytoplasm and are highly networked and elongated (left half of the top cartoon). Mitochondria on stiff ECM are fragmented and clustered near the nucleus (right half of the top cartoon). Stiffness sensitive differential localization of filamin alters mitochondrial localization. Perinuclear enrichment of filamin is responsible for perinuclear clustering of mitochondria (top cartoon). This perinuclear clustering is important for nuclear localization of RUNX2 which primes hMSCs towards osteogenesis on stiff ECM (bottom cartoon). Statistical significance was calculated using two-tailed Mann-Whitney test (**b**) or unpaired t-test with Welch’s correction (**e**). Data are given as median and interquartile range (**b**) or mean ± s.e.m (**e**). P-values are shown in the graphs. White lines denote the cell boundary and cyan ovals depict cell nuclei, both drawn manually using DIC images (for wild-type cells (**a**)) or mApple fluorescence (for transfected cells (**d**)) that were acquired simultaneously with other fluorescence images. Scale bar:10 μm. Soft ECM: 1.2 kPa; Stiff ECM: 90 kPa

## DISCUSSION

Till now, most studies have focused on the effect of matrix stiffening on cytoskeletal architecture, nuclear mechanics, and metabolism. But how these mechanical cues affect the organelle form and function remains mostly unknown. Matrix stiffening and mitochondrial dysregulation both being hallmarks of cancer, ageing, fibrosis, and cardiovascular diseases (45), it is crucial to study the crosstalk between matrix stiffness and mitochondrial form and function. Among the organelles, the mitochondrion is a good starting point not only because of its bioenergetic capabilities but also because it interacts with the cytoskeletal elements and is shown to bear forces. Hence, we chose to focus on the effect of matrix stiffness on mitochondrial morphology and localization. Our study shows that mitochondria in human epithelial cells, MCF-7, and human mesenchymal stem cells are sensitive to mechanical cues. Upon matrix stiffening, they undergo a transition from a homogeneously distributed, filamentous, and highly networked phenotype to a fragmented and less networked phenotype showing perinuclear clustering (Fig. 5f). In this respect, we found that the effect of matrix stiffness is more distinct on mitochondrial localization than on mitochondrial morphology. On the effect of matrix stiffness on mitochondrial morphology, while our results are statistically significant for a p<0.05 and corroborate with the reports of Tharp *et al*. (25), the distribution in morphology is very broad. We believe that this wide distribution may lead to opposing conclusions depending on the context and the study design, which might explain the contradicting results (26, 46). On the contrary, the effect of ECM stiffness on mitochondrial localization is very distinct. To this end, we further show that stiffness-sensitive differential localization of filamin is crucial for determining the subcellular localization of mitochondria. Perinuclear enrichment of filamin on stiff ECM leads to perinuclear mitochondrial clustering (Fig. 3). Going forward, it will be interesting to understand how this perinuclear clustering of fragmented mitochondria is dynamically achieved on matrix stiffening *in vivo*. It is possible that during matrix stiffening due to fibrosis, mitochondria undergo fragmentation to increase ATP activity, are transported to the nucleus in a retrograde fashion by microtubules, and are finally retained near the nucleus by perinuclear actin structures formed by filamin and associated proteins. However, the crosstalk between actin, microtubule and mitochondrial form and localization on soft and stiff ECM needs further investigation. Relevantly, filamin could be a key molecule connecting both mitochondrial form and localization. In fact, given recent reports on stiffness-sensitive regulation of mitochondrial fission proteins (46) and the role of filamin in mitochondrial fission (42), it would be interesting to look at the effect of filamin on localization-dependent or independent dynamics and functions of mitochondria.

Along with localization, we also studied mitochondrial motility by photoactivation and photobleaching. Subsequently, our experiments involving imaging mitochondrial motility revealed that mitochondria on stiff ECM are more mobile (Supp Fig. 3a) and we attribute this behavior to fragmented mitochondria seen on stiff ECM. We also show that on soft ECM, there exist two distinct mitochondrial populations that differ in their motilities (Fig. 2b). PL mitochondria are more mobile than PN mitochondria because of their shorter lengths (minor: major axis) as shown in Supp Fig. 3c. Here, we speculate that the motility difference in PN and PL mitochondrial populations arises due to the difference in the volume constraint of intracellular space near the cell nucleus. The perinuclear region is relatively more organelle-crowded than the peripheral region. The former, therefore, should impose enhanced restrictions to mitochondrial motility as compared to the later. As it appears, the interaction of mitochondria with filamin on stiff ECM abolishes this difference. Beyond physical constraints and molecular interactions, it is also likely that the dynamics of mitochondrial subpopulations is eventually influenced by local subcellular demands. This possibility needs to be further investigated. In fact, distinct mitochondrial populations are also reported in the secretory cells of the salivary gland which have central mitochondria that are mobile and rarely fuse and basolateral mitochondria that are static and frequently fuse (47). Also, pancreatic acinar cells have perinuclear, perigranular and sub-plasmalemmal mitochondria with distinct functions to regulate calcium transport (48). Exploring specific physiological roles of mitochondria in these subcellular niches could therefore provide interesting insights into the spatiotemporal regulation of mitochondria structure and function and add to the under-appreciated heterogeneity in mitochondrial dynamics and positioning.

Finally, while exploring the physiological implication of perinuclear clustering of mitochondria on stiff matrix, we discovered that it is crucial for priming stem cells towards osteogenesis. In this regard, a stiff ECM drives osteogenic differentiation of mesenchymal stem cells in a YAP/TAZ-dependent manner (1, 5). Mitochondrial dynamics and function also play a crucial role in stem cell differentiation (49-52). Upon induction of differentiation, there is a shift from glycolysis to oxidative phosphorylation and mitochondria being at the center of the energy generation process becomes very crucial. While a previous study had just observed perinuclear mitochondria in the early phase of osteogenic induction in MSCs (52), our study reveals a previously unknown role of organelle positioning in priming stem cells towards osteogenesis. We show that stiffness-sensitive perinuclear clustering of mitochondria is important for the nuclear localization of RUNX2, the master regulator of osteogenesis (Fig. 5f). RUNX2 activity has epigenetic regulations, and at the same time, certain mitochondrial metabolites that play a key role in epigenetic modifications, are upregulated upon matrix stiffening (25). Hence, it is possible that perinuclear mitochondrial clustering is crucial not only for successful RUNX2 entry into the nucleus but also for increased mito-nuclear crosstalk in general. This crosstalk is then expected to drive several epigenetic modifications affecting gene transcription. However, whether the perinuclear mitochondria or perinuclear actin cytoskeleton or both facilitate the nuclear entry, sequestration, and activity of RUNX2 and metabolites in the nucleus requires further investigation.

Taken together, considering the recent discoveries of nucleus-associated mitochondria that help in stress resistance and survival (53) and nuclear deformations caused by mitochondrial swelling in cardiomyocytes (54), we speculate that perinuclear mitochondrial clustering in a stiffer microenvironment is an adaptive response that possibly provides an energetic barrier protecting as well as maintaining nuclear architecture and functions.

## MATERIALS AND METHODS

### Cell culture

MCF-7 (NCCS, Pune) cells were cultured in Dulbecco’s modified Eagle’s medium (DMEM) supplemented with GlutaMax (Gibco) with 10% fetal bovine serum (tetracycline-free FBS;Takara Bio) and 10 U ml^-1^ penicillin and 10 μg ml^-1^ streptomycin (Pen-Strep, Invitrogen) in an incubator maintained at 37°C and 5% CO^2^. Human Mesenchymal Stem Cells (hMSCs ; Lonza) were cultured in mesenchymal stem cell basal medium (MSCBM;Lonza) supplemented with mesenchymal cell growth supplement (MCGS; Lonza), L-Glutamine and GA-1000 in an incubator maintained at 37°C and 5% CO^2^.

To establish stable cell lines expressing MitoDendra2, MCF-7 cells were transfected with MitoDendra2 plasmid DNA using Lipofectamine 2000 (Invitrogen). Selection pressure was provided by medium (DMEM-GlutaMax with 10% FBS) containing 400 μg ml^-1^ geneticin (Invitrogen). Stably expressing fluorescent clones were picked using cloning cylinders (Sigma) following fluorescence confirmation. From this suspension, single cells were seeded via serial dilution in a 96-well plate and their growth was monitored for over two weeks. Post this, homogeneously fluorescent colonies derived from single cells were scanned under a fluorescent microscope and sub-cultured into stable cell lines. Subsequent maintenance and passaging of stable cell lines were done in a medium containing 100 μg ml^-1^ geneticin.

Transient transfection with plasmids was done either using Lipofectamine 2000 (Invitrogen) or Xfect (Takara) in a 6-well plate following the manufacturer’s protocol. Post 8-12 h of transfection, cells were trypsinized and seeded onto hydrogel substrates. After at least 24 h in culture, cells were either fixed and immuno-stained or imaged live.

### Hydrogel preparation for compliant ECM

To provide cells with the compliant ECM having different stiffness, polyacrylamide hydrogels coated with collagen-1 were made as described previously (5). 4% (3-Aminopropyl)triethoxysilane(APTES)-treated and 2% glutaraldehyde-activated glass-bottom dishes (Ibidi) were used to cast polyacrylamide (PAA) hydrogel substrates. Hydrogel substrates of varying stiffness with an elastic modulus of 1.2, 4 and 90 kPa were prepared by mixing the desired volume of 40% acrylamide and 2% bisacrylamide as given in Supplementary Table 1). Gel surfaces were functionalized with sulfosuccinimidyl-6-(4′-azido-2′-nitrophenylamino) hexanoate (Sulfo-SANPAH, Thermo Scientific) and covalently coated with 300 μg ml^-1^ collagen-I (Invitrogen) overnight at 4 °C to ensure cell attachment. Cells were seeded onto the gel area and cultured for 24 hours before fixing or live imaging.

### Antibodies, fluorophores, plasmids and other reagents

Source and dilution information for all primary and secondary antibodies and fluorophores are given in Supplementary Table 2. Details of plasmids used in this study are listed in Supplementary Table 3 with their source. Details of all other reagents used in this study are listed in Supplementary Table 4.

### Mitochondrial dyes

For live imaging of mitochondria, cells were incubated with 100 nM Mitotracker Green or Red dye for 15 min prior to imaging.

### Immunofluorescence

Cells were fixed with 4% formaldehyde diluted in 1X phosphate-buffered saline (PBS, pH 7.4) at room temperature (RT) for 15 min, followed by three washes in 1X PBS. The fixed samples were then permeabilized using 0.25% (v/v) Triton X-100 (Sigma) diluted in 1X PBS for 10 min at RT followed by three washes for 5 min each in 1X PBS to remove the detergent. To block non-specific antibody binding, the samples were incubated in 2% BSA in PBST (0.1% v/v Triton X-100 in 1X PBS) at RT for 45 min. The blocking buffer was removed after 45 min, and the primary antibody dilution prepared in the blocking buffer was added to the samples for 60 minutes at RT or at 4°C overnight. After this, samples were washed twice with PBST and thrice with 1X PBS. Samples were then incubated with secondary antibody tagged with Alexa Fluor 488/568/594/647 (same dilution as primary antibody) prepared in blocking buffer, for 60 min at RT in dark. Counter-staining cell nuclei with a DNA-binding dye 4′,6-diamidino-2-phenylindole (DAPI, 1 μg ml^−1^ in PBS) and F-actin with Alexa Fluor dye conjugated phalloidin (Invitrogen) was also done at this step. Finally, samples were washed twice with PBST and thrice with 1X PBS before being imaged using confocal microscopy.

### Confocal microscopy

Immunofluorescence images were acquired using 60X water objective (UPLSAPO W, N.A. = 1.2, Olympus) mounted on an Olympus IX83 inverted microscope equipped with a scanning laser confocal head (Olympus FV3000), Olympus FV31-SW (v2.3.1.198). Time-lapse images of live samples, photoconversion and photo-bleaching studies were done in the same setup using a live-cell chamber at 37°C. HEPES (Gibco) buffer at a final concentration of 50 μM was added to the culture medium to maintain CO2 levels during live imaging.

### Super-resolution microscopy

Images were acquired with a 60X silicone oil objective (<u>UPLSAPO60XS2</u>, N.A=1.3, Olympus) mounted on an Olympus IX83 inverted microscope supplied with Yokogawa CSU-W1 (SoRa Disk) scanner.

### FLIM Imaging and Analysis

For FLIM imaging with Mito Flipper-TR, cells were incubated with 1 μM of the Mito Flipper-TR (Spirochrome) for 15 min. Media conducive for live imaging conditions (fluorobrite DMEM with 10 % FBS, 1 mM Glutamine and 50 μM Hepes buffer) was added to dilute the probe concentration three times and imaging was done in a live cell chamber at 37°C. Imaging was performed using the FV3000 Olympus microscope equipped with a time-correlated single-photon counting module from PicoQuant. A pulsed 485 nm laser (PicoQuant LDH-D-C-485) operated at 20 MHz was used for excitation. The emission was collected through a 600/50 nm bandpass filter, on a gated PMA hybrid 40 detector and a PicoHarp 300 board (PicoQuant). FLIM data was analyzed using the SymPhoTime 64 software (PicoQuant). Rectangular ROIs of comparable areas were drawn manually in perinuclear and peripheral regions of cells. The average fluorescence lifetime was calculated in these ROIs. To determine the fluorescence lifetimes, the fluorescence decay data was fit to a double exponential model after deconvolution for the calculated impulse response function. The values reported in the main text are the average lifetime intensity (tau 1).

### Mitochondrial segmentation and quantification

The workflow and procedures for image processing and thresholding using MitoAnalyser plug-in are described in detail elsewhere (55). In short, 3D image stacks were first acquired and preprocessed using the commands: subtract background, sigma filter plus, enhance local contrast, and gamma correction. To identify mitochondria in the images, “adaptive thresholding” method was used. The settings for thresholding in the Mitochondria Analyser plugin (on FIJI) were adjusted, with ‘scale max slope through stack’ turned on and settings for ‘block size’ and ‘c-value’ adapted per set of images (block size: 2 microns, c:4) with other default settings. The resulting binarized images were post-processed using the despeckle, remove outliers, and fill 3D holes commands. At this stage, each final thresholded image was compared visually to the original image as a quality control check of object identification and segmentation. Image correction, wherever required, was manually done by deselecting some of the pre-processing or post processing commands until the final thresholded image closely resembled the original image. The identified mitochondrial objects were then analyzed in 3D using 3D object counter and “3D particle analyzer” commands to quantify count, volume, surface area, and sphericity. The thresholded objects were also converted into skeletons using “skeletonize (3D)”, and the “analyze skeleton” command was applied to obtain the number of skeletons, number of branches, lengths of branches, and number of branch junctions and end points in the 3D network.

### Mitochondrial motility using photoconversion and photobleaching

MitoDendra2 MCF-7 cells were used for mitochondrial motility studies. MitoDendra2 cells were seeded and cultured on soft and stiff ECM and incubated with 50 nM SiR Actin with 10 μM Verapamil for 12-16 hours prior to imaging. SiR Actin was used to mark cell boundaries that helped in single cell analysis. Photoconversion was done on a rectangular region-of-interest (ROI) in PN and PL regions of MitoDendra2 MCF-7 cell using a 405 nm laser at 4% intensity with a scan speed of 40 μs/pixel. This was immediately followed by LSM imaging of the green and red channels; and mitochondrial motility was subsequently tracked at an interval of 8.5 s for ∼12 mins.

Photobleaching studies were done in MCF-7 cells transiently expressing mCh-KIF5C and Tom20-GFP (RAMP system) and cultured on soft and stiff ECM. These doubly transfected cells were incubated with 50 μM Biotin for an hour to restore mitochondria to its original distribution. 488 nm laser was used at 10% intensity for bleaching a circular region-of-interest (ROI) in PN and PL regions, iterated or looped over three times with a scan speed of 20 μs/pixel. For LSM imaging, the laser power was attenuated to avoid phototoxicity. Photobleaching was done before and after Biotin treatment in the same setup but in different cells. Images were collected before, immediately after, and for ∼1.5 min following the bleaching. All FRAP data were analyzed and mobile fraction was calculated using Stowers plugin in FIJI.

### Analysis of mitochondrial motility using Mitometer

Single cell videos of unconverted mitochondria (green channel) in MitoDendra2 MCF-7 cell line were acquired and fed into the MATLAB plugin of Mitometer for global characterization of mitochondrial motility. Videos of photo converted mitochondria (red channel) in PN and PL regions were cropped out manually and fed into the Mitometer plugin for location specific characterization of mitochondrial motility and morphology. The detailed protocol for running Mitometer is described elsewhere (32). In brief, the average value of all the motility and morphology parameters obtained for each distinct mitochondrial track was calculated and plotted.

### Analysis and quantification of mitochondrial localization

Nucleus and cell body from the respective images were manually segmented and binarized using FIJI. Mitochondria was segmented, thresholded and binarized using the MitoAnalyser plugin in FIJI. Subsequent analysis of mitochondrial and cytoplasmic distribution within a cell as a function of distance from the nucleus was done using a custom MATLAB and python code. Briefly, the centroid of the nuclear mask was determined and considered to be the center of circles for subsequent radial distribution analysis. For all the non-zero pixels of the mitochondrial mask, the distance from the nuclear centroid was calculated and written into a “csv” file. After subtracting the nucleus mark from the cell body mask, the distance of all the non-zero pixels of the cell body mask from the nuclear centroid was calculated and written into another “csv” file. The minimum distance obtained here is considered as the nuclear radius. The “perinuclear region” was defined as the region that falls within 5 microns from the nuclear radius. These output files from the MATLAB code were then used for further analysis in Python. Inbuilt functions of the Python seaborn library were used for radial distribution analysis. Normalized frequency distribution of non-zero mitochondrial and cell body pixels with distance from the nucleus was determined using the “distplot” function. In the histogram of non-zero mitochondrial pixels, the area within the perinuclear region is the fraction of perinuclear mitochondrial content. All distances in pixels were converted to microns prior to analysis.

### Image Analysis

To determine perinuclear filamin, ROIs were traced out manually using the selection brush tool (fixed at 20-pixel width) in FIJI. Cell-nuclei frame was synced with the filamin frame and used as a reference for tracing the perinuclear region. Whole cell boundaries were manually traced using DIC or phalloidin frames. Total intensity values were taken for perinuclear and whole cell regions per cell. The fraction of perinuclear filamin localization per cell was quantified as the ratio of total intensity of the perinuclear region to the total intensity of the whole cell. All fluorescence images were brightness-adjusted and denoised uniformly throughout the whole image for representation purposes only. Denoising was done using the PureDenoise tool in FIJI.

### Statistics and Reproducibility

Statistical analyses were carried out in GraphPad Prism 9 (Version 9.2.0). Tests done for statistical significance and respective p-values are mentioned in all the figures. *p*-Values > 0.05 were considered to be statistically not significant. No statistical methods were used to set the sample size. Quantification was done using data from at least two independent biological replicates. All the experiments with representative images were repeated at least twice.

## Supporting information

Combined Supplementary Information

## AUTHOR CONTRIBUTIONS

P.D. and T.D. formulated the project. P.D. performed the majority of the experiments and analysis. B.T developed and wrote a code for robust analysis of mitochondrial localization.

S.R. exclusively performed and analyzed FLIM-based membrane tension and contributed to other experiments as well. P.D. and T.D. wrote the manuscript. All authors agreed on the manuscript as in the submitted version.

## ACKNOWLEDGEMENT

We thank Manish Jaiswal and Kalyaneswar Mandal for critical discussion and Shilpa P. Pothapragada for assistance in data analysis. We thank Vaishnavi Ananthnarayan and Narendrakumar Ramanan for generously providing us with plasmids. T.D. is a DBT/Wellcome Trust India Alliance intermediate fellow and partner group leader of the Max Planck Society (MPG), Germany. Authors sincerely acknowledge generous funding from the Human Frontier Science Program (HFSP) Research Grant (RGP0007/2022), DBT/Wellcome Trust India Alliance (Ref. No. IA/I/17/1/503095), and intramural funds at TIFR Hyderabad from the Department of Atomic Energy (DAE), India, under Project Identification Number RTI 4007, towards supporting this project.

