## Supplementary material for "Matrix stiffening promotes perinuclear clustering of mitochondria": Combined Supplementary Information

Daga *et al.*

This document contains:

1. Supplementary Table 1: Composition of compliant polyacrylamide gels with different stiffness.
2. Supplementary Table 2: Antibody/Fluorophore information
3. Supplementary Table 3: Plasmid information
4. Supplementary Table 4: Reagent information
5. Supplementary Video Legends
6. Supplementary Figure Legends
7. Supplementary Figures 1-6

**Supplementary Table 1. Composition of compliant polyacrylamide gels with different stiffness.**

| Stiffness (Young's Modulus) | 1.2 kPa | 4 kPa | 90 kPa |
|---|---|---|---|
| milli-Q $H_2O$ (µl) | 832 | 819 | 554 |
| 40% acrylamide (µl) | 137 | 125 | 300 |
| 2% bis-acrylamide (µl) | 25 | 50 | 140 |
| 10% APS (µl) | 5.75 | 5.75 | 5.75 |
| TEMED (µl) | 0.25 | 0.25 | 0.25 |

**Supplementary Table 2. Antibody/Fluorophore information**

| Antibodies/Fluorophores | Catalog No. | Manufacturer | Dilution/ Working concentration |
|---|---|---|---|
| Mouse anti-ATP5A (monoclonal) | ab14748; clone 15H4C4 | Abcam | 1:100 |
| Rabbit anti-RUNX2 (monoclonal) | 12556S; clone D1L7F | CST | 1:1000 |
| Mouse anti-Filamin A (monoclonal) | F6682; clone PM6/317 | Sigma | 1:100 |
| DAPI (4′,6-diamidino-2-phenylindole) | D1306 | Invitrogen | 1 µg $ml^{-1}$ |
| Anti-rabbit IgG (H+L), F(ab')2 Fragment (Alexa Fluor-647 Conjugate) | 4414S | CST | Same as per primary antibody dilution |
| Anti-rabbit IgG(H+L), F(ab') Fragment (Alexa Fluor 594 Conjugate) | 8889 | CST | |
| Alexa Fluor 647, Goat anti-Mouse IgG (H+L) Cross-Adsorbed Secondary Antibody | A21236 | Invitrogen | |
| Alexa Fluor 568, Goat anti-Mouse IgG (H+L) Cross-Adsorbed Secondary Antibody | A11031 | Invitrogen | |
| Alexa Fluor 488, Goat anti-Mouse IgG (H+L) Cross-Adsorbed Secondary Antibody | A-11001 | Invitrogen | |
| Alexa Fluor 488, Goat anti-Rabbit IgG (H+L) Cross-Adsorbed Secondary Antibody | A-11008 | Invitrogen | |
| Mito Flipper-TR | CY-SC023 | Cytoskeleton | 1 µM |
| SiR-Actin | CY-SC006 | Cytoskeleton | 50 nM |
| Alexa Fluor 647 Phalloidin | 8940 | CST | 1:40 |
| Mitotracker Green | 9074S | CST | 100 nM |
| Mitotracker Red CMXRos | 9082S | CST | 100 nM |

**Supplementary Table 3: Plasmid information**

| Plasmid | Source | Identifier | Comment |
|---|---|---|---|
| mApple-FilaminA-N-9 | Addgene | 54901 | Filamin overexpression |
| mApple-dnFLNA | This work | DOI: 10.1073/pnas.1104 21 1108 | Human FLNA Repeat [19- 22]-HA cloned into mApple C1 |
| Strep-KifC1*-mCherry | Addgene | 120169 | KIF-C1 based retrograde motor tagged with Streptavidin and mCherry |
| TOM20*-SBP-GFP | Addgene | 120173 | Mitochondrial OMM protein Tom20 tagged with SBP and GFP |
| Mito-Dendra 2 | Vaishnavi Ananthanarayanan | Addgene: 55796 | Photo-convertible mitochondrial marker |
| Vinculin T (pCAG-C1-Vcl-T) | Narendrakumar Ramanan | DOI: 10.1007/s00018-021-03879-7 | Vinculin Tail domain overexpression |
| Vinculin T12 (pCAG-C1-Vcl-T12) | Narendrakumar Ramanan | DOI: 10.1007/s00018-021-03879-7 | Constitutively active Vinculin |

**Supplementary Table 4: Reagent information**

| Reagents | Catalog No. | Manufacturer |
|---|---|---|
| Dulbecco's Modified Eagle Medium (DMEM) GlutaMAX | 10569010 | Invitrogen |
| Fetal Bovine Serum (Tet-free) | 631101 | Takara |
| Penicillin-Streptomycin (10,000 U/mL) | 15140122 | Thermo |
| Trypsin-EDTA (0.25%) | 25200056 | Invitrogen |
| Opti-MEM I | 31985062 | Invitrogen |
| Human Mesenchymal Stem Cells (hMSCs) | PT-2501 | Lonza |
| Mesenchymal Stem Cell Basal Medium (MSCBM) | PT-3238 | Lonza |
| Mesenchymal Cell Growth Supplement (MCGS) | PT-4106E | Lonza |
| L-Glutamine | PT-4107E | Lonza |
| GA-1000 | PT-4504E | Lonza |
| Fluorobrite DMEM | A1896701 | Invitrogen |
| HEPES | 15630 | Gibco |
| L-Glutamine Solution | G7513 | Sigma |
| Geneticin | 10131035 | Invitrogen |
| Formaldehyde | 28908 | Thermo |
| Bovine Serum Albumin | MB083 | Himedia |
| Triton X 100 | T8787 | Sigma |
| Xfect Transfection Reagent | 631317 | Takara |
| Lipofectamine 2000 | 11668019 | Invitrogen |
| Sodium Hydroxide Solution | 415413 | Sigma |
| (3-Aminopropyl) trimethoxy silane 97% | 281778 | Sigma |

| | | |
|---|---|---|
| Glutaraldehyde | G6257 | Sigma |
| 40 % acrylamide | A4058 | Sigma |
| N,N'-Methylenebisacrylamide solution (2 % in water) | M1533 | Sigma |
| Ammonium Persulfate | A3678 | Sigma |
| N,N,N′,N′-Tetramethylethylenediamine | T7024 | Sigma |
| Sulfo-SANPAH | 803332 | Sigma |
| Collagen 1 (rat tail) | A1048301 | Invitrogen |
| Biotin | B4639 | Sigma |
| Verapamil | CY-SC006 | Cytoskeleton |

**Supplementary Video Legends:**

**Supplementary Video 1**:
Mitochondria on soft ECM (4 kPa) have subpopulations that differ in their motilities.
Left: Time lapse video showing mitochondrial dynamics in unconverted mitochondria (green) and photo converted mitochondria (red) of a MitoDendra2 MCF-7 cell cultured on soft ECM. Right: Time lapse video showing mitochondrial dynamics of photo converted mitochondria (in black). Photoconversion was carried out at PN and PL regions (yellow boxes) and the dynamics of photo converted mitochondria was tracked at an interval of 15s for ~23 mins. Diffusion of photoconverted molecules was faster in PL mitochondria when compared to PN mitochondria highlighting differences in mitochondrial motility. Montage is shown in Fig 2a. Movie is shown at 5 frames per second. PN: perinuclear, PL: peripheral. Scale bar :10 µm.

**Supplementary Video 2:**
Mitochondria on stiff ECM (90 kPa) have subpopulations that have comparable motilities.
Left: Time lapse video showing mitochondrial dynamics in unconverted mitochondria (green) and photo converted mitochondria (red) of a MitoDendra2 MCF-7 cell cultured on stiff ECM. Right: Time lapse video showing mitochondrial dynamics of photo converted mitochondria (in black). Photoconversion was carried out at PN and PL regions (yellow boxes) and the dynamics of photo converted mitochondria was tracked at an interval of 6s for 6 mins and then at an interval of 15s for ~26 mins. Diffusion of photoconverted molecules was faster in PN mitochondria when compared to PL mitochondria indicating networked mitochondria near the nucleus and highlighting location specific differences in mitochondrial morphology. However, motilities of PN

and PL mitochondria were comparable. Montage is shown in Supp Fig 3b. Movie is shown at 5 frames per second. PN: perinuclear, PL: peripheral. Scale bar: 10 µm.

**Supplementary Video 3:**
Perinuclear clustering of mitochondria on soft ECM (4 kPa) abolishes differences in mitochondrial motility. Perinuclear clustering of mitochondria in MCF-7 cells was achieved by transient transfection of mCh-KIF5C and Tom20-GFP cultured on soft ECM (4 kPa) without the addition of biotin (- Biotin) (see Supp Fig 3d). Photo-bleaching was carried out at T= 00:04s in PN and PL regions (white circles) and the recovery of fluorescence was monitored for 1 min 40 s. The fluorescence recovery in PN and PL mitochondria were comparable. Montage and FRAP curves shown in Fig 2c (top panel) highlight that perinuclear mitochondrial clustering abolishes differences in mitochondrial motility. Movie is shown at 5 frames per second.
PN: perinuclear, PL: peripheral. Scale bar: 10 µm.

**Supplementary Video 4**:
Reversing perinuclear clustering of mitochondria to its original homogenous distribution on soft ECM (4 kPa) reestablishes mitochondrial populations with different motilities.
Perinuclear clustering of mitochondria in MCF-7 cells achieved by transient transfection of mCh-KIF5C and Tom20-GFP cultured on soft ECM (4 kPa) was reversed upon addition of biotin and mitochondria were reverted back to their original homogenous distribution within the cell. After an hour of incubation with biotin (+ Biotin), photo-bleaching was carried out at T=00:04s in PN and PL regions (white circles) and the recovery of fluorescence was monitored for 1 min 40 s. The fluorescence recovery in PL mitochondria was higher than PN mitochondria reiterating our observation that soft ECM generates mitochondrial populations that have different motilities. Montage and FRAP curves shown in Fig 2c (bottom panel). Movie is shown at 5 frames per second. PN: perinuclear, PL: peripheral. Scale bar: 10 µm.

**Supplementary Figure Legends:**

**Supplementary Figure 1:**

**a**. Representative fluorescence images showing filamentous mitochondria on soft ECM (top panel) and toroidal mitochondria on stiff ECM (bottom panel). Mitochondria (yellow) in MCF-7 cells were visualized live using Mitotracker Green dye. Scale bar: 10 μm.

**b**. Scatter dot plots showing network properties of mitochondria imaged under live conditions: Branches, Branch Junctions, Branch Endpoints and Total Branch Length per mitochondria in MCF-7 cells cultured on soft (4 kPa) and stiff (90 kPa) ECM. n=24 FOVs (4 kPa), 25 FOVs (90 kPa). Each FOV (field of view) contains 10-15 cells.

**c**. Scatter dot plots showing weighted sphericity of mitochondria per cell and network properties namely branches, branch junctions, branch endpoints and total branch length per mitochondria in MCF-7 cells cultured on soft (4 kPa) and stiff (90 kPa) ECM and imaged under fixed conditions after immunostaining. n=6 FOVs (4 kPa), 6 FOVs (90 kPa). Each FOV contains 10-15 cells.

**d**. Scatter dot plots showing weighted sphericity of mitochondria per cell and network properties namely branches, branch junctions, branch endpoints and total branch length per mitochondria in wildtype (WT) and Vcl-T12 transfected cells on soft (4 kPa) ECM and wildtype (WT) and Vcl-T transfected cells on stiff (90 kPa) ECM. n= 18 (WT, 4 kPa), 15 (Vcl-T12, 4 kPa), 16 (WT, 90 kPa), 18 (Vcl-T, 90 kPa) cells.

Statistical significance was calculated using two-tailed Mann-Whitney test (**b, c**) or Kruskal-Wallis nonparametric test with uncorrected Dunn's test (**d**). Data are given as median and interquartile range (**b, c, d**) taken over two independent replicates. P-values are shown in the graphs. Soft ECM: 4 kPa; Stiff ECM: 90 kPa

**Supplementary Figure 2:**

**a**. Immunofluorescence confocal images of MCF-7 cells cultured on soft (top panel) and stiff (bottom panel) ECM under untransfected (wildtype) and transfected conditions showing mitochondria stained with ATP5α (in yellow), nuclei stained with DAPI (in cyan) and actin stained with phalloidin (in gray). From left to right: Merged image of MCF-7 wildtype cells showing nuclei in cyan and mitochondria in yellow followed by actin in grayscale highlighting distinct actin stress fibres on stiff ECM, merged image of MCF-7 cells transfected with Vcl-T12 (magenta) showing nuclei in cyan and mitochondria in yellow followed by actin in grayscale showing actin stress fibres on soft ECM, merged image of MCF-7 cells transfected with Vcl-T (magenta) showing nuclei in cyan and mitochondria in yellow followed by actin in grayscale showing diffused actin structures on stiff ECM.

**b-c.** Binarized images of cells shown in Figure 1f. Binarized images of cell body, nucleus and mitochondria in MCF-7 cells transfected with Vcl-T12 (**b**) or Vcl-T (**c**) and cultured on soft ECM (4 kPa, top panel) and stiff ECM (90 kPa, bottom panel). From left to right: Binarized images showing area of a single cell (in black) followed by nuclear area (in black) and mitochondria

positive pixels (in white), graphs showing distribution of cytoplasm (blue) and mitochondria (orange) plotted as a function of distance from the nucleus (localization distribution function). The nuclear, perinuclear, and cytoplasmic zones are indicated on the graphs.
Scale bar: 10 μm. Soft ECM: 4 kPa; Stiff ECM: 90 kPa

**Supplementary Figure 3:**
**a**. Scatter dot plots showing motility metrics: speed (μm $s^{-1}$), displacement (μm) and velocity (μm $s^{-1}$) of mitochondrial tracks on soft (4 kPa) and stiff (90 kPa) ECM. n=1168 (4 kPa), 1586 (90 kPa) distinct mitochondrial tracks in MitoDendra2 MCF-7 cells (unconverted state i.e., GFP fluorescence).
**b**. Montage showing mitochondrial motility at perinuclear (PN) and peripheral (PL) regions of MitoDendra2 expressing MCF-7 cells cultured on stiff ECM (90 kPa). Yellow boxes represent ROIs where Dendra2 expressing mitochondria were photoconverted (t=00:00) and subsequently the diffusion of the photoconverted molecules was tracked for 20 mins (also see Supp Video 2).
**c**. Scatter dot plots showing minor: major axis (refers to minor and major axis of an ellipse. High value of this ratio indicates more circular mitochondria) of photo converted mitochondria in PN and PL regions of MitoDendra2 MCF-7 cells on soft (4 kPa) and stiff (90 kPa) ECM. n= 546 (soft PN), 594 (soft PL), 469 (stiff PN), 675 (stiff PL) distinct mitochondrial tracks.
**d**. Confocal image showing perinuclear clustering of mitochondria (in yellow) in an MCF-7 cell transiently transfected with mCh-KIF5C (magenta) and Tom20-GFP (yellow) and cultured on soft ECM (4 kPa). Photo-bleaching and subsequent recovery and FRAP analysis (shown in Fig 2c, top panel; Supp Video 3) was carried out in this doubly transfected cell. White line denotes the cell boundary drawn manually using mCherry fluorescence image. Scale bar :10 μm.
**e**. Scatter dot plots showing mobile fractions calculated in PN and PL regions of MCF-7 cells transfected with mCh-KIF5C and Tom20-GFP and cultured on soft ECM (4 kPa, left graph) and stiff ECM (90 kPa, right graph) in the absence (- Biotin) or presence (+ Biotin) of biotin. n= 5 (Soft PN, - Biotin), 5 (Soft PL, -Biotin), 7 (Soft PN, + Biotin), 7 (Soft PL, + Biotin), 5 (Stiff PN, - Biotin), 5 (Stiff PL, -Biotin), 4 (Stiff PN, + Biotin), 4 (Stiff PL, + Biotin) ROIs from cells. (5 cells on soft ECM without biotin, 7 cells on soft ECM with biotin, 5 cells on stiff ECM without biotin, 4 cells on stiff ECM with biotin).
Statistical significance was calculated using unpaired nonparametric Kolmogorov Smirnov test (**a**) or Kruskal-Wallis nonparametric test with uncorrected Dunn's test (**c, e**). Data are given as median and interquartile range (**a, c, e**). P-values are shown in the graphs. The Y-axes of plots showing, speed, displacement and velocity are on the "log 10" scale. Other plots are on the linear scale.

**Supplementary Figure 4:**
**a**. Immunofluorescence images of MCF-7 cells stained for filamin cultured on soft (top panel) and stiff (bottom panel) ECM followed by representative grayscale images with traced-out ROIs.
Perinuclear regions and cell boundaries traced manually for quantifying perinuclear fraction of filamin.
**b-d**. Binarized images of cells shown in Figure 3d. Binarized images of cell body, nucleus, and mitochondria in wildtype MCF-7 cells (**b**) or cells transfected with mAppFLNA (**c**) or dnFLNA (**d**) and cultured on soft ECM (top panel) and stiff ECM (bottom panel). From left to right:

Binarized images showing area of a single cell (in black) followed by nuclear area (in black) and mitochondria positive pixels (in white). The corresponding graphs are shown in Fig 3e.
Scale bar: 10 μm. Soft ECM: 4 kPa; Stiff ECM: 90 kPa

**Supplementary Figure 5:**
**a.** Confocal images showing co-localization of Mito Flipper- TR (in green) and Mitotracker Red (in red) probes in MCF-7 cells. Scale bar: 10 μm.

**Supplementary Figure 6:**
**a**. Scatter dot plot showing network properties namely branches, total branch length, branch junctions and branch endpoints per mitochondria in hMSCs cultured on soft (1.2 kPa) and stiff (90 kPa) ECM. n=18 FOVs (1.2 kPa), 15 FOVs (90 kPa) taken over two independent replicates. Each FOV (field of view) contains 3-5 cells. Statistical significance was calculated using two-tailed Mann-Whitney test. Data are given as median and interquartile range. P-values are shown in the graphs.
**b**. Binarized images of cells shown in Figure 5d. Binarized images of cell body, nucleus, and mitochondria in wildtype and mAppFLNA transfected hMSCs cultured on soft ECM (top panels) and in wildtype and dnFLNA transfected hMSCs cultured on stiff ECM (bottom panels). From left to right: Binarized images showing area of a single cell (in black) followed by nuclear area (in black) and mitochondria positive pixels (in white). The corresponding graphs are shown in Fig 5c.
**c**. Immunofluorescence images of hMSCs cultured for 1 day on soft ECM (left image) and stiff ECM (right image) and stained with DAPI (blue) and RUNX2 (green). RUNX2 puncta are only detectable in nuclei on stiff ECM.
**d**. Immunofluorescence images of hMSCs cultured on stiff ECM under untransfected (wildtype, left panel) and transfected (dnFLNA, right panel) conditions and stained with DAPI, RUNX2 and ATP5α. From top to bottom: Merged images showing nuclei in cyan and mitochondria in yellow, grayscale images of mitochondria. Magnified views of the red boxes are shown in Fig 5e. White lines represent the cell boundary drawn manually using DIC images that were acquired simultaneously with other fluorescence images. Scale bar: 10 μm. Soft ECM: 1.2 kPa; Stiff ECM: 90 kPa

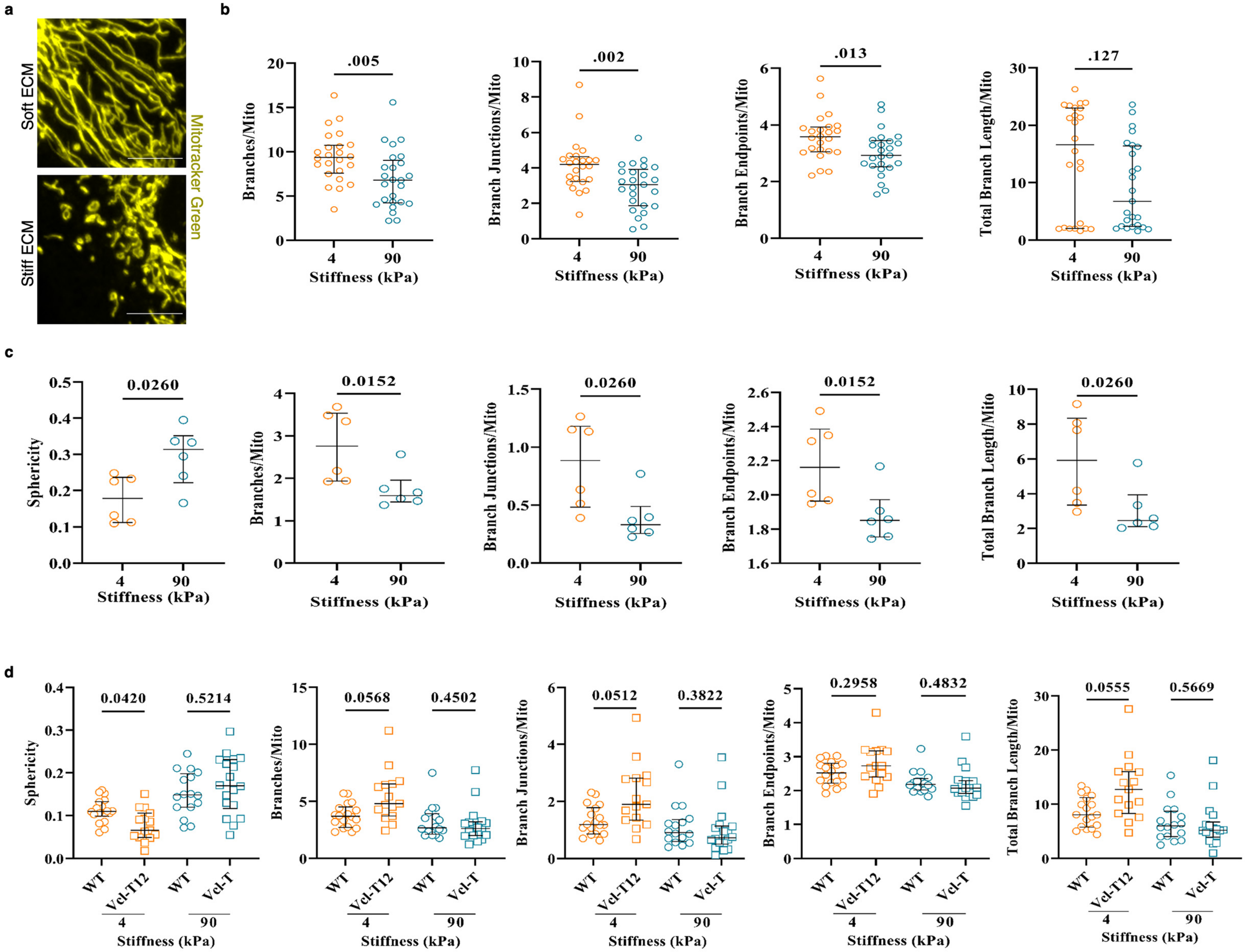


Supplementary Figure 1

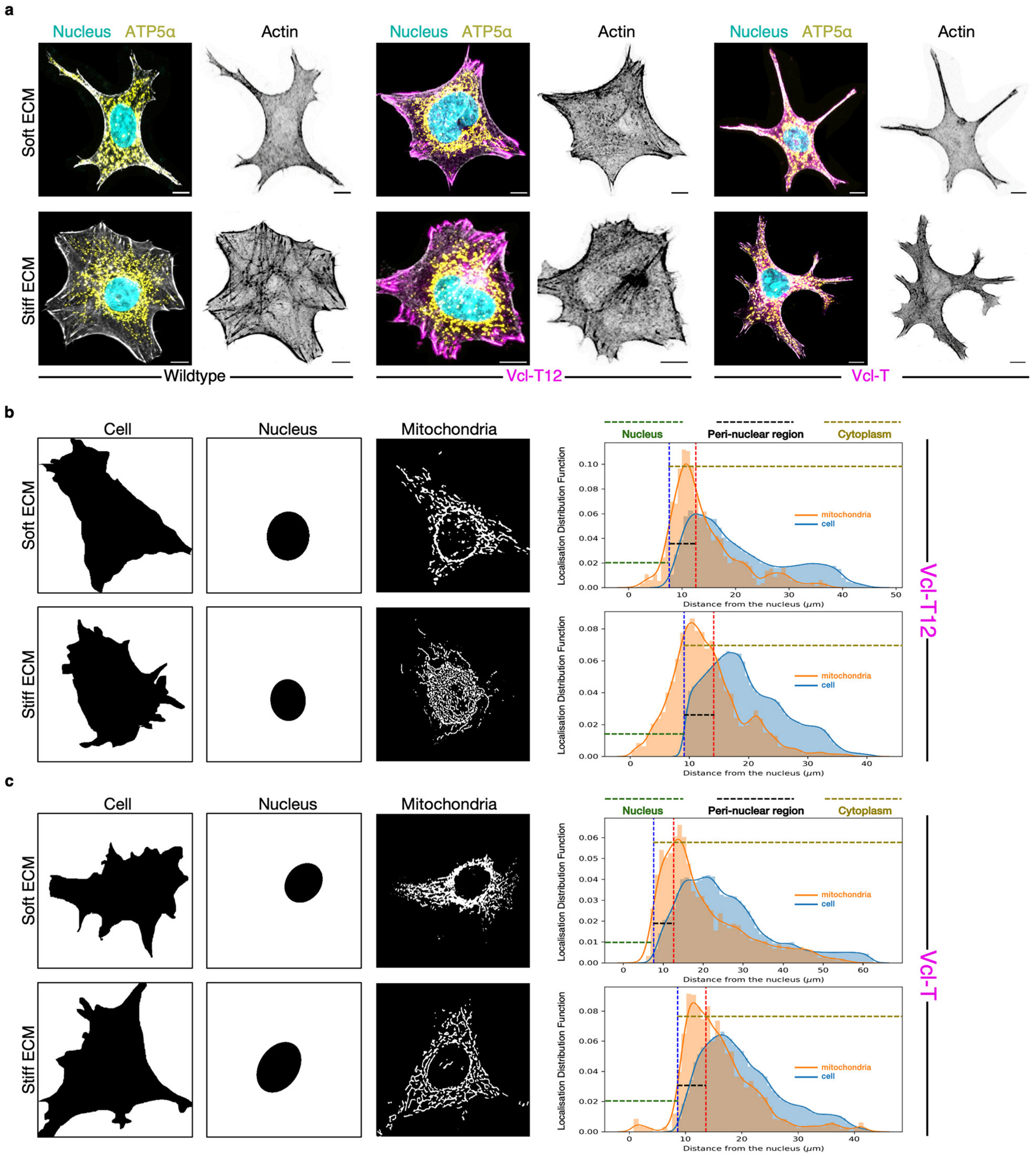


**Supplementary Figure 2**

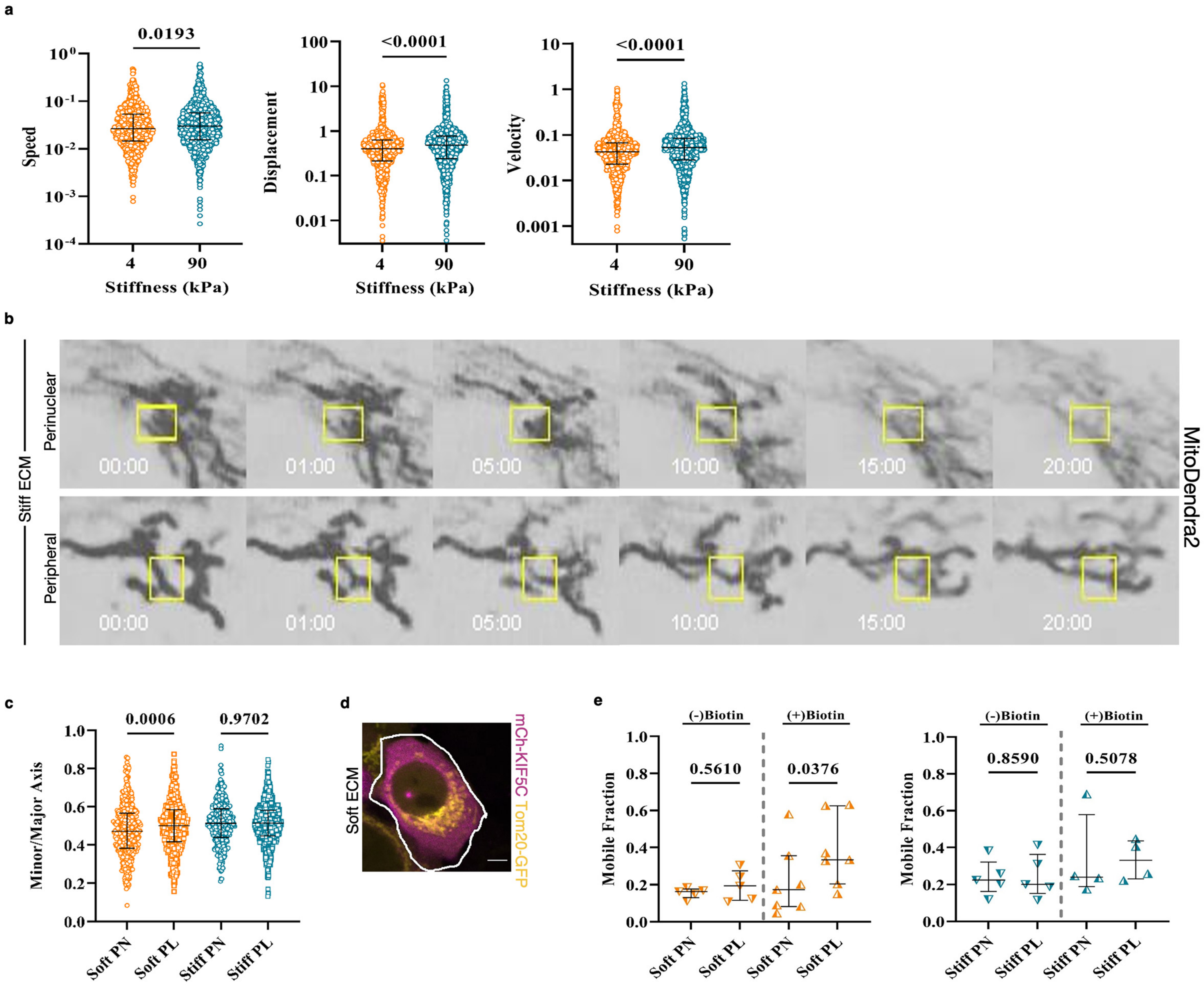


**Supplementary Figure 3**

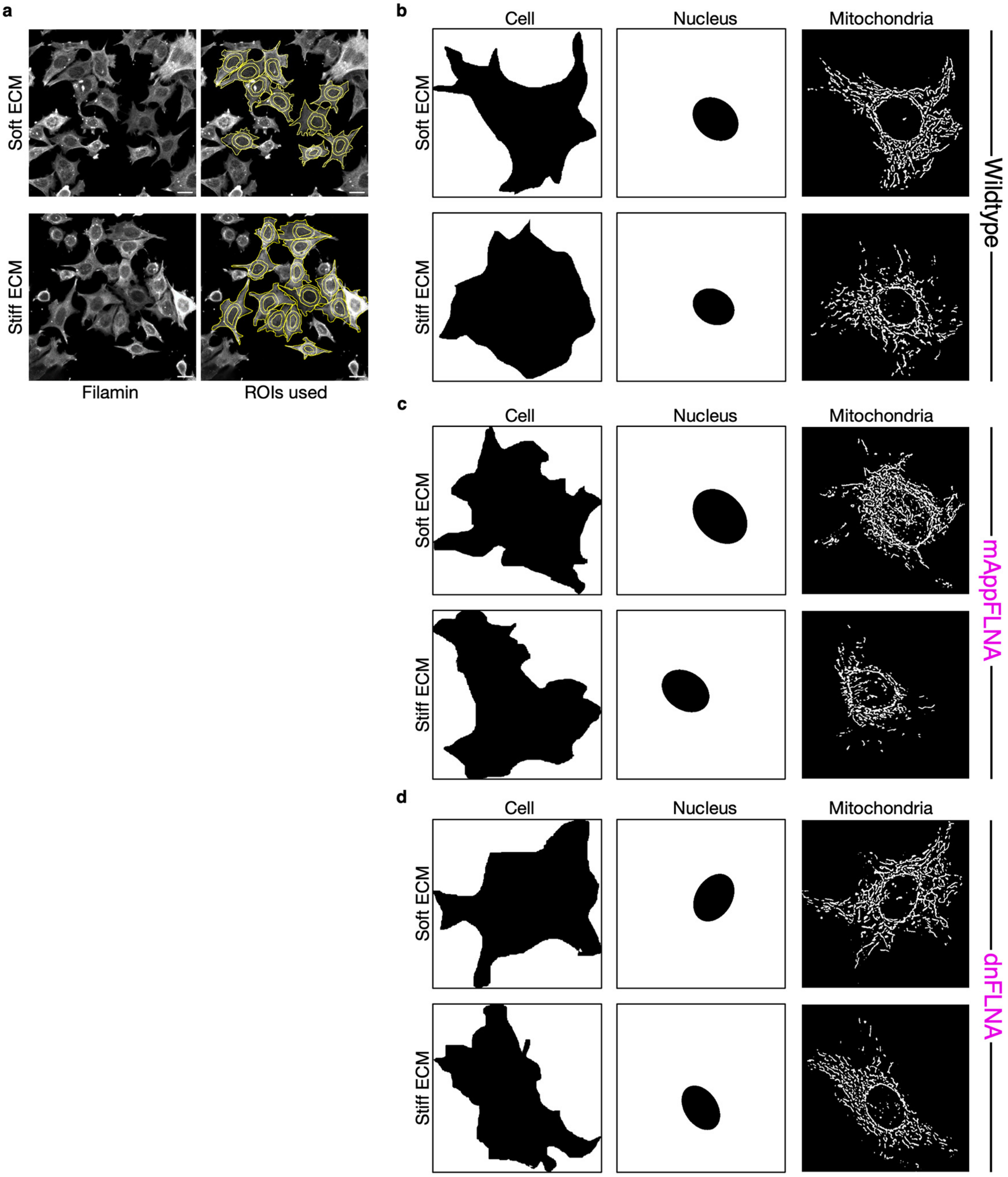


**Supplementary Figure 4**

a

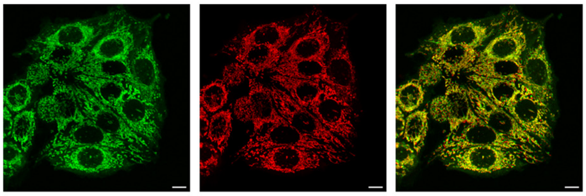

Mito Flipper-TR Mitotracker Red Merge

### Supplementary Figure 5

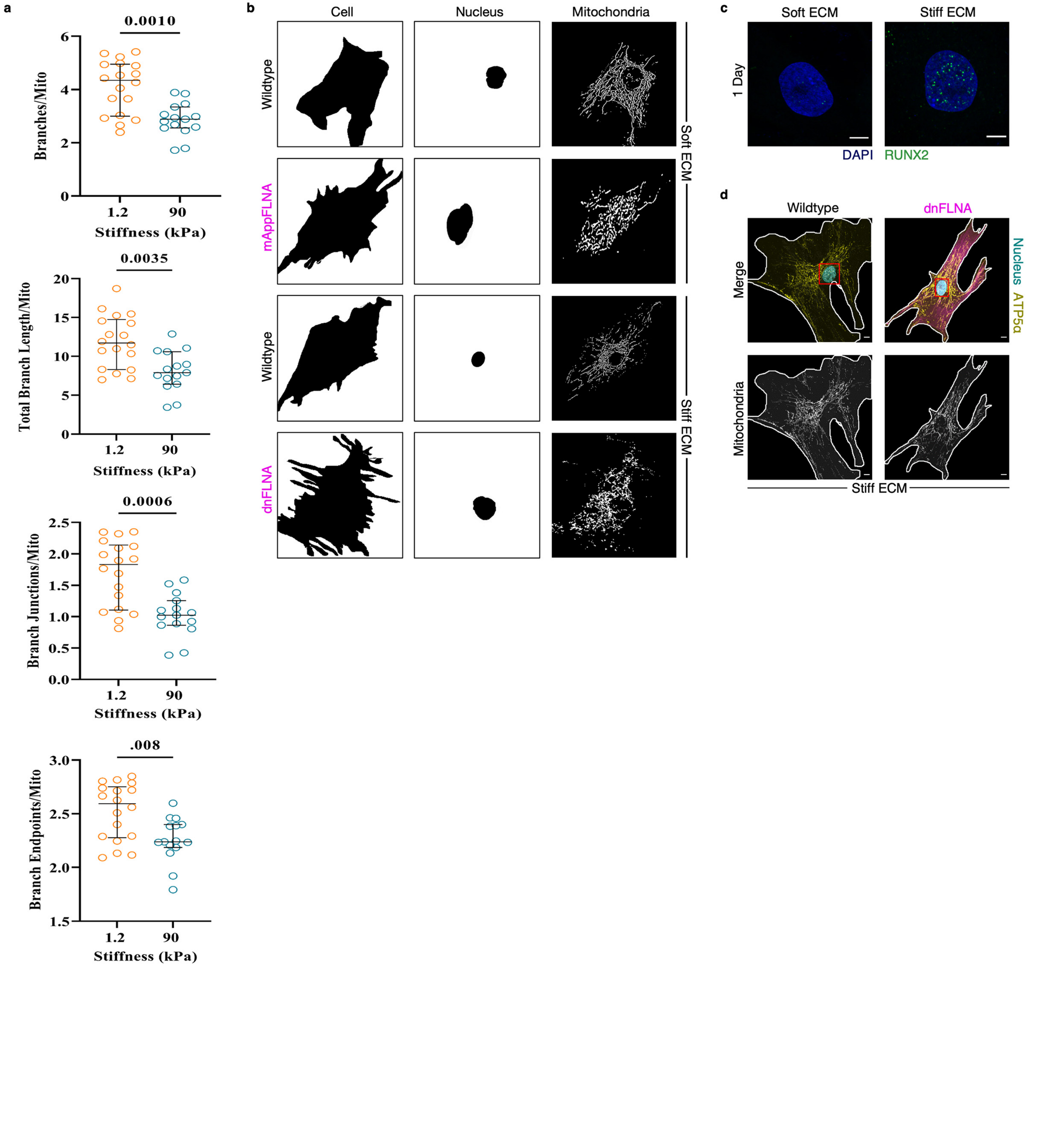


**Supplementary Figure 6**
